# Topological segregation of functional networks increases in developing brains

**DOI:** 10.1101/378562

**Authors:** Wei He, Paul F. Sowman, Jon Brock, Andrew C. Etchell, Cornelis J. Stam, Arjan Hillebrand

## Abstract

A growing literature conceptualises human brain development from a network perspective, but it remains unknown how functional brain networks are refined during the preschool years. The extant literature diverges in its characterisation of functional network development, with little agreement between haemodynamic- and electrophysiology-based measures. In children aged from 4 to 12 years, as well as adults, age appropriate magnetoencephalography was used to estimate unbiased network topology, using minimum spanning tree (MST) constructed from phase synchrony between beamformer-reconstructed time-series. During childhood, network topology becomes increasingly segregated, while cortical regions decrease in centrality. We propose a heuristic MST model, in which a clear developmental trajectory for the emergence of complex brain networks is delineated. Our results resolve topological reorganisation of functional networks across temporal and special scales in youth and fill a gap in the literature regarding neurophysiological mechanisms of functional brain maturation during the preschool years.

## Introduction

Modern network science has revealed that normal brain networks exhibit fundamental properties of three canonical network extremes - a *random network* (***Erdös and Rényi, 1959***), a locally connected and highly *ordered* (regular) network (***Mulder, 1992***), and a *scale-free network* with a small number of highly connected nodes (so-called “hubs”, ***Barabasi and Albert 1999***). Adult brain networks also display hierarchical modularity (***Meunier et al., 2009; Stam, 2014; Wig, 2017***), in which modules that include regions from the default mode, fronto-parietal, parieto-temporal, or subcortical networks support specific cognitive functions (***Bullmore et al., 2009; Fornito et al., 2011; Power et al., 2011***). A heuristic model of complex brain networks has been proposed (***Stam and van Straaten, 2012***) to characterise the properties of real brain networks in an abstract “network space” defined by the four network models (i.e., regular, random, scale-free, and hierarchical modular networks). This heuristic model of “network space” suggests that the hierarchical modular network is an “attractor” for healthy brain networks and the other three extreme networks are “attractors” for different stages or patterns of brain diseases (***Stam and van Straaten, 2012; Stam, 2014***).

Despite the robust and reproducible description of adult brain networks, there is relatively scant data regarding the maturation of brain networks. Such data can be acquired non-invasively using magnetic resonance imaging (MRI) or electrophysiological techniques (such as magnetoencephalography/MEG and electroencephalography/EEG). Studies using MRI-based measurements have demonstrated that both functional and structural brain networks become more segregated during childhood (e.g., functional MRI: ***Fair etal. 2009; Guetal. 2015; Supekar et al. 2009***; structural MRI: ***Huang et al. 2015***; and diffusion-weighted imaging:***Baum et al. 2017***). Such development allows for an ongoing balance between the *integration* of converging information from distributed brain regions, and at the same time the *segregation* of divergent specialised information streams (***Fair et al., 2009; Grayson and Fair, 2017; Richmond et al., 2016; Rubinov and Sporns, 2010***). However, most studies to date have only focused on children older than 6 years or younger than 3 years of age (***Grayson and Fair, 2017***), leaving the preschool years of childhood (between 3 and 6 years of age) understudied – a knowledge gap that has been termed “the missing neurobiology of cognitive development” (***Poldrack, 2010***).

Furthermore, there is little agreement between MRI- and electrophysiology-based network descriptions. Correspondence between functional MRI and electrophysiological measures of functional brain networks (***Brookes et al., 2011***) implies that changes in functional MRI network organisation should be, at least partially, preserved in higher temporally-resolved electrophysiological investigations (***Grayson and Fair, 2017***). It follows then, that electrophysiological networks are expected to become increasingly segregated during childhood development. However, prior EEG studies have reported conflicting results, which include increasing segregation (***Boersma et al., 2011, 2013; Janssen et al., 2017; Toth et al., 2017***), decreasing segregation (***Smit et al., 2016; Bathelt et al., 2013; Miskovic et al., 2015***), or no changes with age (***Schafer et al., 2014***). Discrepancy between developmental MRI-and electrophysiology-based network findings has been difficult to reconcile, partly due to the different spatial scales that functional networks have been examined at (sensor-level in most EEG versus cortical-level in fMRI studies). Modern whole-head magnetoencephalography (MEG) allows for sophisticated spatial filtering techniques to accurately (varying from sub-millimetre to a few centimetres) reconstruct millisecond electrophysiological time series across the cortex (***Hillebrand et al., 2005; Troebinger et al., 2014; Barratt et al., 2018***), and thus MEG is a critical tool in the quest to resolve these discrepancies.

To better understand how the topology of functional brain networks develops over the whole period of childhood, we used MEG to collect resting-state electrophysiological signals from children whose ages spanned 4 to 12 years, as well as from adults. Importantly, we utilised a paediatric MEG system with a child-sized helmet for data collection in children aged under 6 years (***He et al., 2014; Johnson et al., 2010***). We hypothesised that, based on the heuristic model of complex brain networks, the healthy brain develops from a more random and integrated structure towards a configuration that offers a balance between network integration and segregation during normative development (***Stam, 2014***). Specifically, we predicted that: (1) functional networks become more segregated, shifting from a centralised network topology to a de-centralised configuration (***Boersma et al., 2013; Toth et al., 2017***); (2) individual brain regions become more diverse in their connectedness, i.e., centrality of brain regions increases for hubs (e.g., regions in the default mode and the fronto-parietal areas), but decreases in non-hub regions (e.g., regions in the primary visual and auditory areas).

## Results

We applied an atlas-based beamforming approach (***Hillebrand et al., 2012***) to reconstruct time series of neuronal activity recorded using a child-customised 125-channel whole-head gradiometer MEG system optimised for children aged around 5 years (5 year-olds (Y.O.), N = 10, 5.4 ± 1.1 years, 5 males). We used a 160-channel whole-head gradiometer MEG system for children aged around 10 years (10 year-olds (Y.O.), N = 14, 9.8 ± 1.5 years, 12 males) and adults (N = 24, 40.6 ± 17.4 years, 16 males). Functional connectivity between the 80 regions of interest (ROIs; 78 cortical ROIs and bilateral hippocampi) in the automated anatomical labelling (AAL; ***Tzourio-Mazoyer et al. 2002***) atlas was estimated using the phase lag index (PLI). Averaged PLI was computed between a region and all 79 other regions, resulting in a single estimation of functional connectivity per participant. There were no significant PLI differences between the three age groups for any of the 5 frequency bands (delta: 0.5–4 Hz, theta: 4–8 Hz, alpha: 8–13 Hz, beta: 13–30 Hz, and low gamma: 30–48 Hz).

Subsequently, we reconstructed the minimum spanning tree (MST; ***Figure 1; Kruskal 1956; Wang et al. 2008***), so that the topology of functional networks could be characterised and compared without biases that are inherent in conventional graph theoretical approaches (***Stam, 2014; Tewarie et al., 2015***). The MST is a sub-network that contains the strongest connections within a weighted network without forming cycles or loops; it provides an unbiased reconstruction of the core of a network, making it possible to create a unique backbone or empirical reference network (e.g., for large datasets such as the human brain connectome project; ***van Dellen et al. 2018***). Moreover, MST parameters are sensitive to alterations in the topology of brain networks at the functional- (e.g., ***Boersma et al. 2013; de Bie et al. 2012; Janssen et al. 2017)*** and structural-level (e.g., ***Otte et al.2015; van Dellen et al. 2018***), and importantly, can be interpreted along the lines of conventional graph theoretical measures (***Tewarie et al., 2016***).

**Figure 1.**
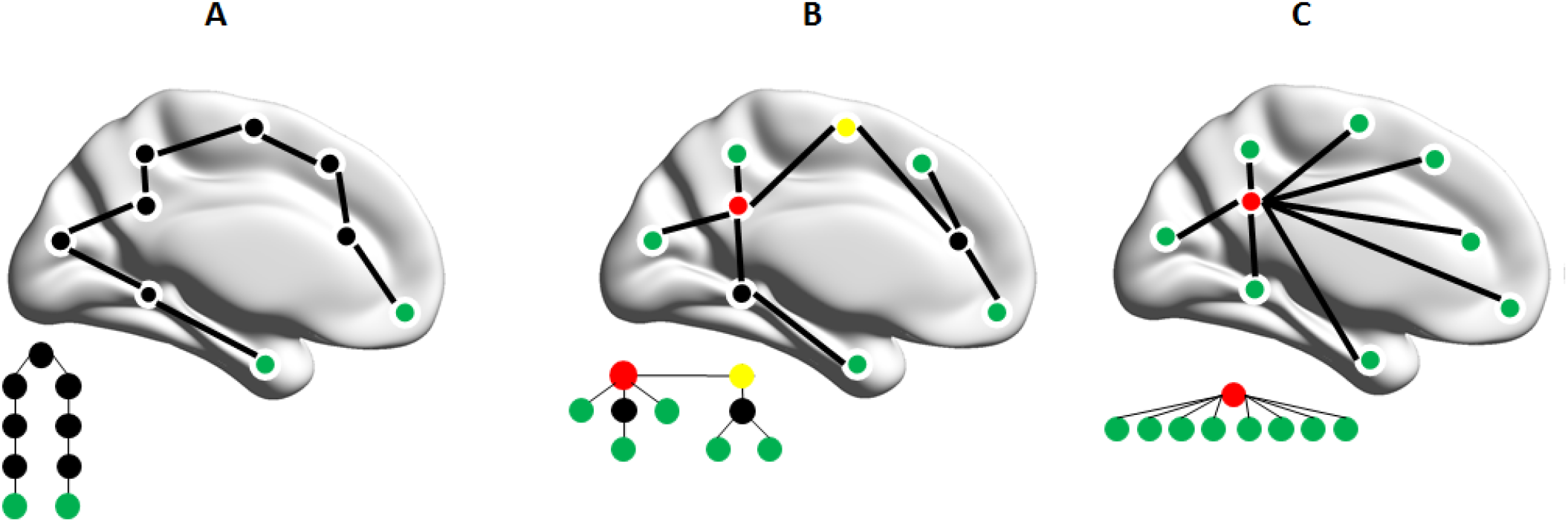
Minimum spanning tree (MST) topology and hierarchy of three representative tree models. Top panel: (A) a line-like tree, and (C) a star-like tree. (B) an intermediate configuration between the two extremes. Nodes are indicated by circles, and links by connecting lines. Green nodes are leaves, which have a ***Degree*** (i.e., number of links to neighbouring nodes) of 1; red nodes are hubs that have the highest ***Degree*** and ***Betweenness Centrality*** (i.e., the fraction of the smallest number of links between any two nodes in a network that pass through a node); the yellow node and the red node in B, have the lowest ***Eccentricity*** (i.e., the largest number of links required for a node reaching any other node in a network). The ***Diameter*** in B is 5 (i.e., the longest distance between any two nodes in a network). The three lower graphs are the same trees as those overlayed on the template brains above but represented in a way that illustrates that trees with more leaves have fewer layers (nodes with the lowest *Eccentricity* are placed on top). Network A requires many steps for an individual node, especially a leaf node in green, to connect to other nodes (low *integration* and high *segregation*). The steps required for nodes to connect with each other are fewer in C but the central hub/red node is considered ‘overloaded’ (high *integration* but low *segregation*). The network between these extremes - network B - represents a hierarchical tree, which offers a balance between information *integration* and *segregation*.

### Topological segregation of the large-scale functional networks

We first sought to understand whether the topology of the functional networks become more segregated during childhood development. To this end, we calculated 5 global MST measures for each participant: Diameter, Leaf Fraction, Tree Hierarchy, Degree Correlation, and Kappa. Small Diameter and high Leaf Fraction are characteristic for a highly integrated topology such as a star-like network (A in *Figure 1*), whereas large Diameter and low Leaf Fraction are representative of a more segregated topology or line-like network (C in *Figure 1*). An optimal MST topology, requiring a small Diameter without overloading central nodes, is quantified by Tree Hierarchy (***Boersma et al., 2013; Tewarie et al., 2015***). Such a network topology also tends to have larger Degree Correlation and Kappa, suggesting it is resilient against random damage (***Barrat et al., 2008; Van Mieghem et al., 2010***).

The 5 global MST measures were significantly different across all 5 frequency bands when comparing children (as a whole group) to adults: Kappa, Leaf Fraction, and Tree Hierarchy were higher, whereas Degree Correlation and Diameter were lower, in the children (***Figure 2***). These frequency-independent effects were all highly significant (*p* < 0.001) when contrasting 5 Y.O. with the other two age groups, but less so when comparing 10 Y.O. with adults. The 10 Y.O was adult like for most global MST topological measures, apart from larger Leaf Fraction in the delta (*p* = 0.036) and beta (*p* = 0.041) bands, larger Kappa (*p* = 0.017) and Leaf Fraction (p = 0.036) in the theta band, and smaller Diameter (*p* = 0.023) but larger Leaf Fraction (*p* = 0.038) and Tree Hierarchy (*p* = 0.007) in the alpha band. Overall, the MST topology becomes more line-like and segregated across all frequency bands with increasing age (***Figure 3***).

**Figure 2.**
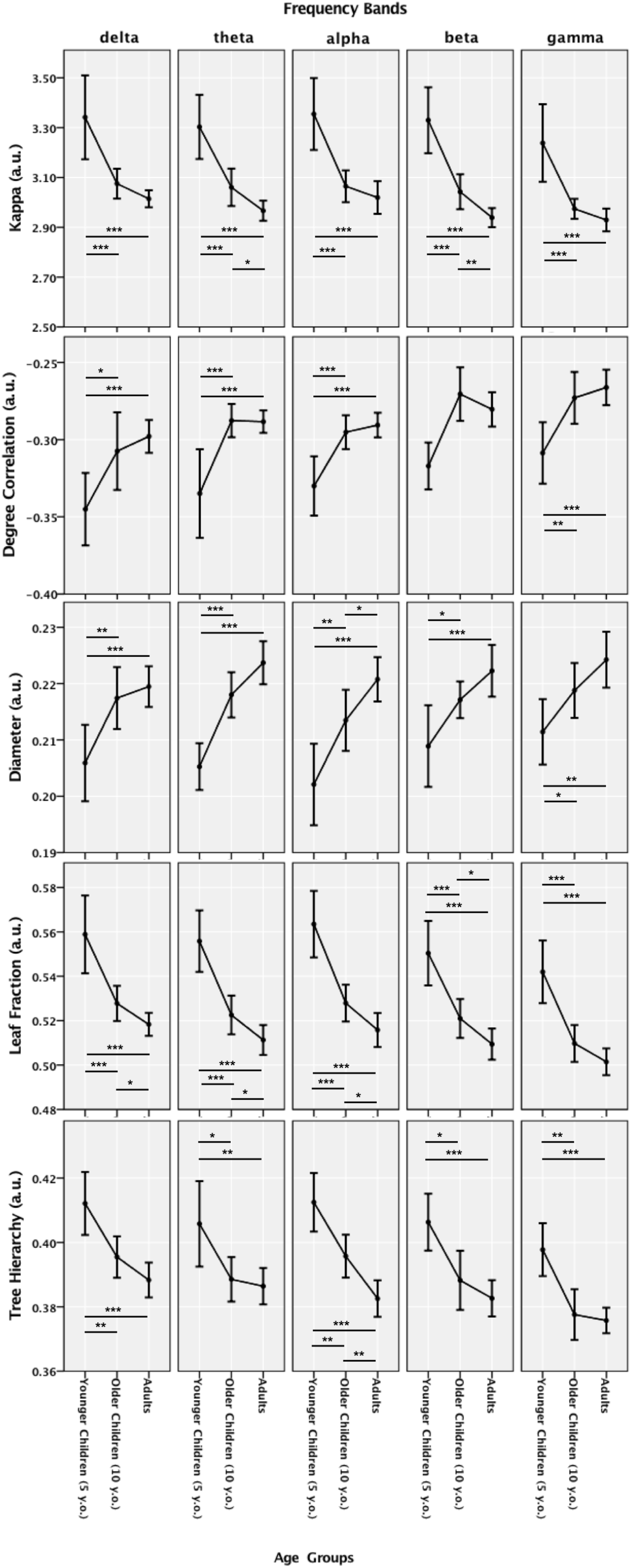
Minimum spanning tree (MST) global metrics estimated from individual phase lag index adjacency matrices in the delta (0.5–4 Hz), theta (4–8 Hz), alpha (8–13 Hz), beta (13–30 Hz), and low gamma (30–48 Hz) bands for three age groups (5 year-olds (Y.O.), 10 year-olds (Y.O.), and Adults). Error bars depict 95% confidence intervals estimated using bootstrapping with 1000 random iterations. * indicates statistically significant group differences (p < 0.05, 50000 random permutations), ** for *p* < 0.01, and *** for *p* < 0.001.

**Figure 3.**
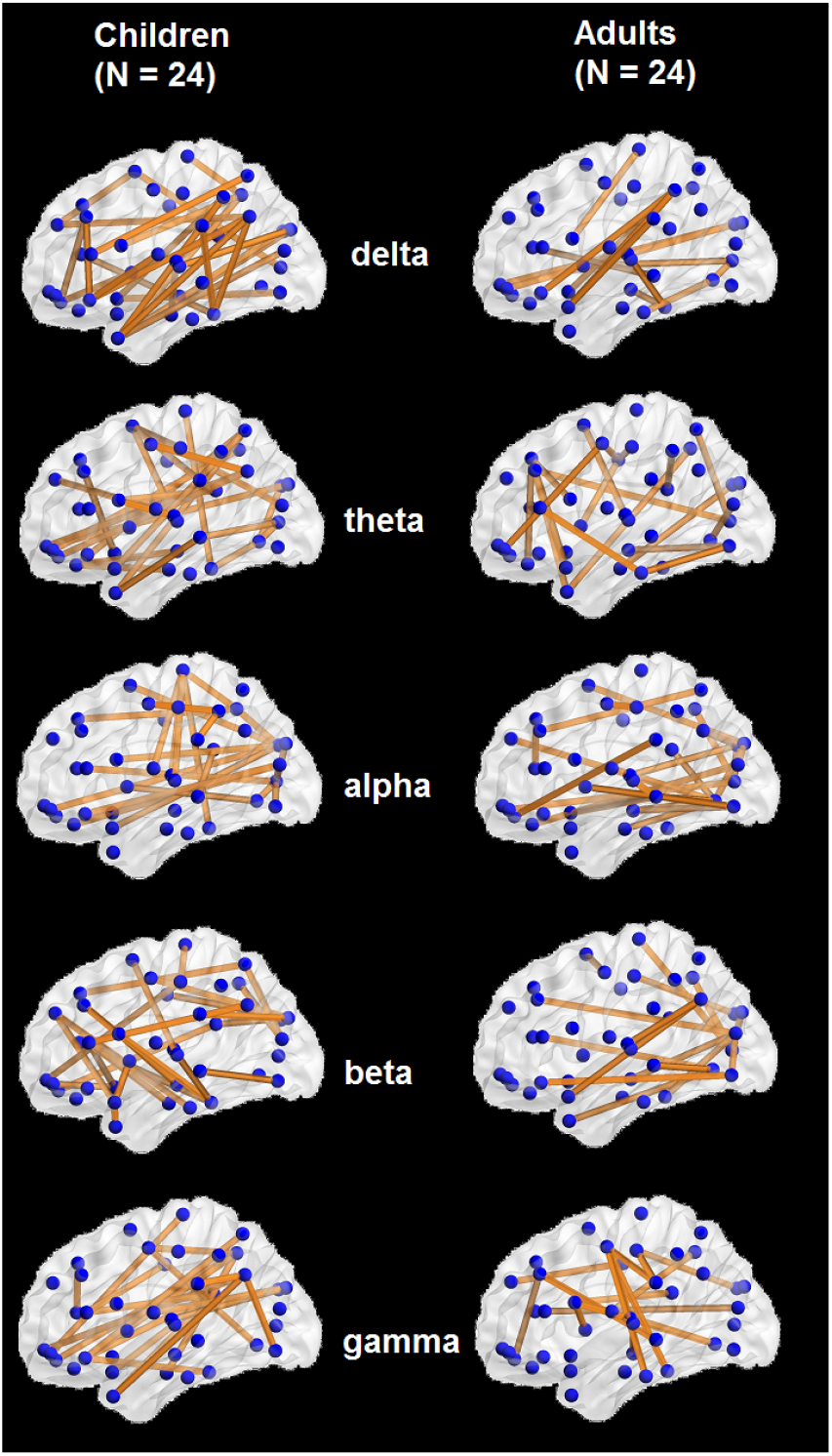
Minimum spanning trees (MSTs) for adults (N = 24) and children (N =24) in five frequency bands (delta: 0.5-4 Hz, theta: 4-8 Hz, alpha: 8-13 Hz, beta: 13-30 Hz, and low gamma: 30-48 Hz), displayed on a template brain with blue dots depicting nodes and yellow lines depicting functional connections. The MSTs depicted are estimated from averaged phase lag index adjacency matrices from adults (right panel) and children (left panel) for illustrative purposes only. The alpha-mediated MST in adults has fewer leaves and a more line-like topology (with central nodes in occipital regions) than the MST in children. This observation agrees with the statistical comparisons between age groups when the MST metrics were based on the un-averaged adjacency matrices in ***Figure 2.***

### Regional de-centralisation correlates with increasing topological segregation

Having established that the network topology is more segregated in adults than in children, we next investigated the centrality of brain regions. We calculated 3 nodal MST measures for each of the 80 regions in every participant: Degree, Betweenness Centrality, and Eccentricity. Larger Degree and Betweenness Centrality, but smaller Eccentricity characterise regions (or so-called “hubs”) that play a central role in the network. We found that, even though there were no significant group differences for the Degree and Betweenness Centrality, the Eccentricity showed significant increases from children (as a whole group) to adults, and from 5 Y.O. to adults in particular. The group differences for the Eccentricity, illustrated in ***Figure 4***, show pervasive changes in Eccentricity over the cortex (the full results are shown in Tables 1-5 in ***Appendix 1***).

**Figure 4.**
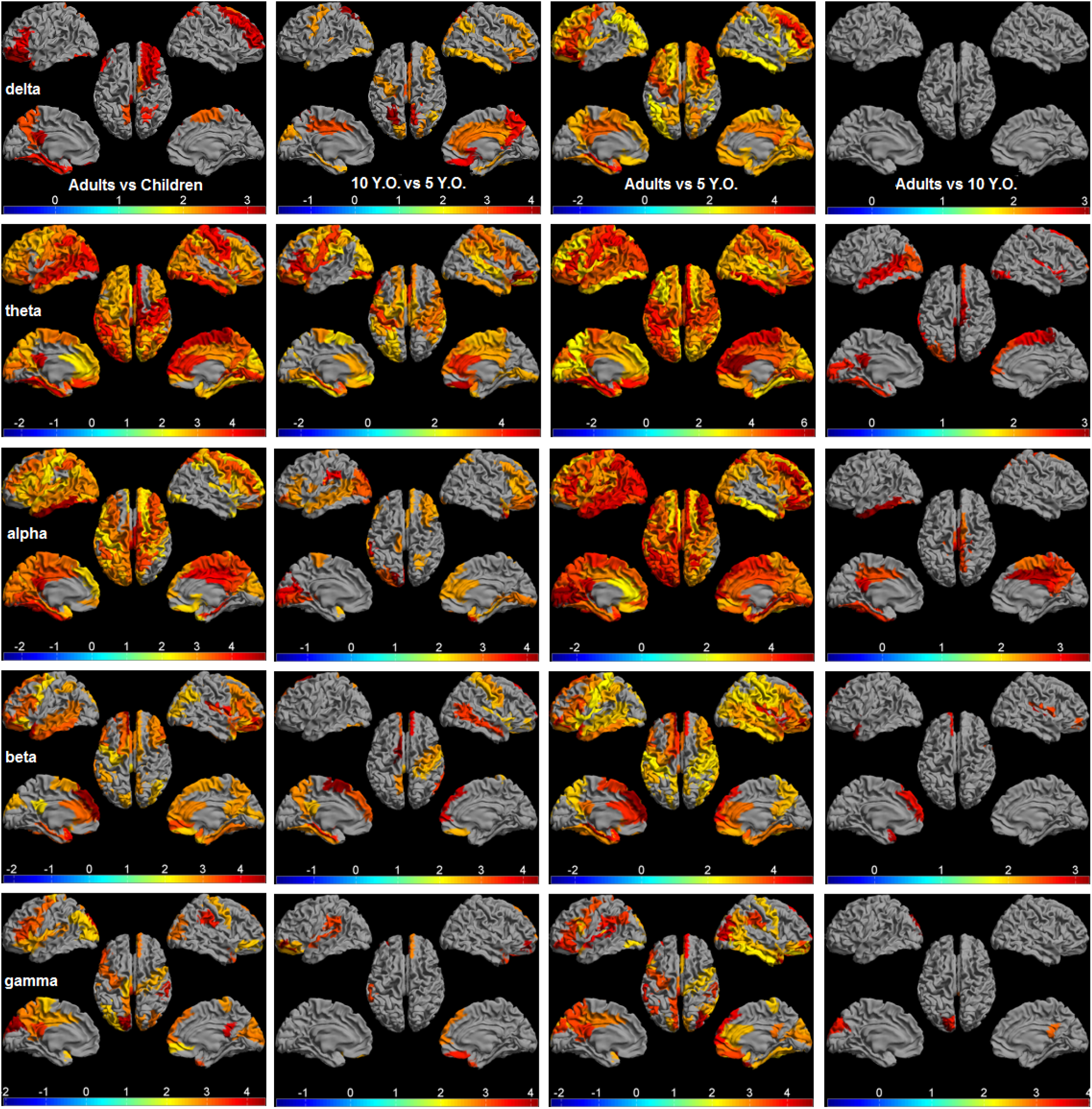
Significant differences in the minimum spanning tree (MST) Eccentricity displayed as a color-coded map on the parcellated template brain, viewed from, in clockwise order, the left, top, right, right midline, and left midline. From left to right, pairwise differences (t-value, p < 0.05, FDR-corrected for 3 nodal MST measures x 80 ROIs) between adults and children, 10 Y.O. and 5 Y.O., adults and 5 Y.O., as well as adults and 10 Y.O., are shown for all five frequency bands (delta: 0.5-4 Hz, theta: 4-8 Hz, alpha: 8-13 Hz, beta: 13-30 Hz, and low gamma: 30-48 Hz).

When contrasting adults and 5 Y.O.:

- all 80 ROIs showed larger theta band Eccentricity in adults;
- in alpha, beta, and delta mediated MSTs, most of the nodes showing larger Eccentricity were in fronto-parietal areas, followed by the nodes normally assigned to the default mode and parieto-temporal areas, and in hippocampal and occipital areas;
- about half of the nodes in the default mode, parieto-temporal, and the occipital areas showed larger Eccentricity in gamma mediated MSTs.

When comparing adults and 10 Y.O.:

- most of the nodes showing larger Eccentricity were in the default mode, occipital, parieto-temporal, and fronto-parietal areas in alpha band mediated MSTs;
- nodes from the default mode, parieto-temporal and occipital areas showed larger Eccentricity in the theta mediated MSTs;
- nodes from the fronto-parietal, parieto-temporal, and hippocampal areas, as well as the nodes from the default mode, showed larger Eccentricity in the beta mediated MSTs;
- only nodes from occipital area and the default mode area showed larger Eccentricity in the gamma mediated MSTs;
- no Eccentricity differences were found in the delta mediated MSTs.

When contrasting adults to children (as a whole group), and 5 Y.O. to the other two age groups, the group differences in Eccentricity exhibited a similar pattern, namely that a larger Eccentricity was found mostly in nodes from the fronto-parietal area, followed by those from default mode, parieto-temporal, occipital and hippocampal areas in delta-to-gamma mediated MSTs.

## Discussion

Capitalising on several novel approaches, we demonstrate in this cross-sectional MEG study that the topology of functional brain networks becomes segregated during childhood development. Increasing topological segregation is associated with increasing regional Eccentricity across the cortex, indicating that most brain regions become functionally specialised and less central in the network. Specifically, the reorganisation of network topology has the same profile across all frequency bands and is not routed via a few hub regions. Importantly, all topological network differences are highly significant between the preschool children/5 Y.O. and older age groups (i.e., older children/10 Y.O. and adults), suggesting that the preschool years present a unique and important period of network maturation. These converging results on topological network changes inform a heuristic MST model from which normal development during childhood can be characterised.

The delineation of large-scale functional brain networks in adults has confirmed a number of hypotheses regarding the degradation of network function in aging and disease (***Stam, 2014***). However, the small number of developmental studies that have examined electrophysiological networks have produced heterogeneous results. Furthermore, these results do not align well with MRI-based haemodynamic imaging data. Critically, we resolved these discrepancies by utilising several technical and methodological advances: (1) age-appropriate MEG systems that are insensitive to age-related physiological and anatomical changes in biological tissues (e.g., bone thickness and density of the skull; ***Smith et al. 2012***); (2) source-level functional connectivity estimation to facilitate interpretation of our results in an anatomical context, and to effectively mitigate spurious connectivity/network results inherent in sensor-level analyses (***Antiqueira et al., 2010; Lai et al., 2017***); (3) leakage insensitive connectivity estimation using PLI, which effectively ignores spurious connectivity due to field spread (***Dominguez et al., 2007***) and volume conduction/signal leakage (***Lai et al., 2017; Schoffelen and Gross, 2009; Stam et al., 2007***); (4) lastly, MST for unbiased network comparisons between different age groups (***Tewarie et al., 2015; Van Mieghem et al., 2010***).

Leveraging data across multiple frequency bands in anatomical space, we demonstrate that the topology of electrophysiological networks becomes increasingly segregated during childhood, in line with MRI-based findings (***Baum et al., 2017; Fair et al., 2009; Gu et al., 2015; Huang et al., 2015***). The smaller Diameter and larger Leaf Fraction in children compared to adults indicates that the topology of the functional brain networks becomes segregated via a transition from a star-like (centralised) configuration toward a more line-like (de-centralised) configuration during development. Such network topological change has been found in infants right after birth (***Toth et al., 2017***) and continues up to 18 years of age (***Boersma et al., 2011***). In addition, the observed larger Kappa in children compared to adults suggests a movement away from a scale-free network. This finding seems to be at odds with findings from most adult studies, which indicate that the mature brain network is approximately “scale-free” (***Sporns, 2013***). However, Kappa is not strictly tied to “scale-freeness”, but rather is a measure for the homogeneity of the degree distribution in the MST (especially in the case of small networks; ***Jinhui et al. 2009***). Moreover, scale-freeness is a relative measure, and depends on the reference model that the experimental model is compared to (***Stam and van Straaten, 2012***). Thus, the adult brain may still be scale-free, although less so than brain networks in children. In accordance with the decreased scale-freeness of adult networks, the increase in Eccentricity found in a distributed set of brain regions across all frequency bands suggests that during development most brain regions, including hubs become less central, in order to prevent hub overloading, as well as to reduce vulnerability to targeted attacks (***Stam et al., 2009***). Together, decreasing nodal centrality possibly reflects a protective mechanism during normative brain development, since disturbances and insults to hub regions can produce lifelong changes in neurological and mental functioning (***Crossley et al., 2014; DeSalvo et al., 2014; Stam et al., 2009; Tewarie et al., 2014; Yu et al., 2017***). Lastly, the smaller Tree Hierarchy found in adults is less straightforward to understand here, as a decrease in network hierarchy is often observed in clinical groups (***Stam and van Straaten, 2012***). Tree Hierarchy is a composite MST measure that takes into account several aspects of the MST, namely the maximum Betweenness Centrality and the number of leafs (***Stam, 2014***). Given that Betweenness Centrality and Degree did not differ between children and adults, the observed decrease in Tree Hierarchy, in our data, is likely to be driven by a decrease in Leaf Fraction. A more straightforward quantification of network hierarchy, other than Tree Hierarhcy, in complex network neuroscience is warranted though. Nevertheless, the present data point to a balance between network integration and segregation (i.e., a network topology that becomes increasingly segregated) with locally specialised regions, during childhood development.

Most network differences in the current study are frequency-independent, suggesting that similar network constraints manifest themselves across different physiological architectures (***Barry et al., 2004; Bathelt et al., 2013; Murias et al., 2007***). All global MST changes in our study share the same profile across the five frequency bands between age groups. Although the specific distributed regions that showed centrality differences varied across frequency bands, there were also some frequency invariant differences: the largest number of regions that exhibited between group Eccentricity differences was found in theta and alpha mediated MSTs; regions in the frontoparietal and default mode areas displayed the largest differences across all frequency bands. This seems to contradict some frequency-specific network findings reported in lower frequency bands in previous developmental EEG studies (***Boersma et al., 2011; Miskovic et al., 2015; Srinivasan, 1999***). These inconsistencies may be ascribed to differences between cohorts (e.g., age-profiles) and methodological differences (e.g., the use of weighted versus unweighted graphs, use of different thresholds, and/or the normalisation of networks/graphs via random surrogates; ***van Wijk et al. 2010***). Nevertheless, MST analysis used in our study effectively addresses methodological limitations such as biased estimates of network topology and biased network comparisons (***Tewarie et al., 2015***).

Furthermore, there is now a growing understanding that conventional graph theoretical metrics (such as the clustering coefficient and shortest path length) do not fully account for fundamental properties of brain networks, and the small-world model is often used inappropriately in the field of neuroscience (***Papo et al., 2016***). Therefore, we propose here a heuristic MST model space to better capture the trajectory of changes in functional brain networks underlying normative brain development (***Figure 5***). Within this MST model space, current findings suggest a clear developmental trajectory of brain networks along the right axis, suggesting a balance between integration and segregation in topology. An adequate delineation of different trajectories of topological changes in abnormal development, which may be a more useful biomarker than the absolute values (***Wolff and Piven, 2014***), can also be provided by this network space. For instance, MST networks were found to be more star-like in ADHD children compared to age-matched typical children *Janssen et al., 2017)* - a pattern that fits with a shift towards the lower-right corner of the network space. Such a trend indicates a delay in brain maturation for ADHD children. In contrast, MST networks become more line-like in children with dyslexia compared to typically developing children (***Fraga Gonzalez et al., 2016***) - a transition to the lower-left corner of the network space. This pattern indicates an alternative developmental trajectory along the horizontal axis for brain networks in dyslexia, veering from the typical developmental trajectory along the right axis. Our model space suggests that the normal adult brain that emerges during development is a special composite that combines optimal network integration and segregation, degree diversity, and hierarchy. Moreover, distinct pathological trajectories in adults, if projecting the normal adult brain onto the horizontal axis, could also be represented in this model space: a more de-centralised line-like MST was found in patients with early relapsing remitting multiple sclerosis (***Tewarie et al., 2014***) and Alzheimer’s disease (***Yu et al., 2016***), suggesting that networks in these diseases move towards the lower-left corner (more segregated); a more centralised star-like MST was observed in fronto-temporal dementia (***Yu et al., 2016***), indicating an opposite trend towards the lower-right corner (more integrated).

**Figure 5.**
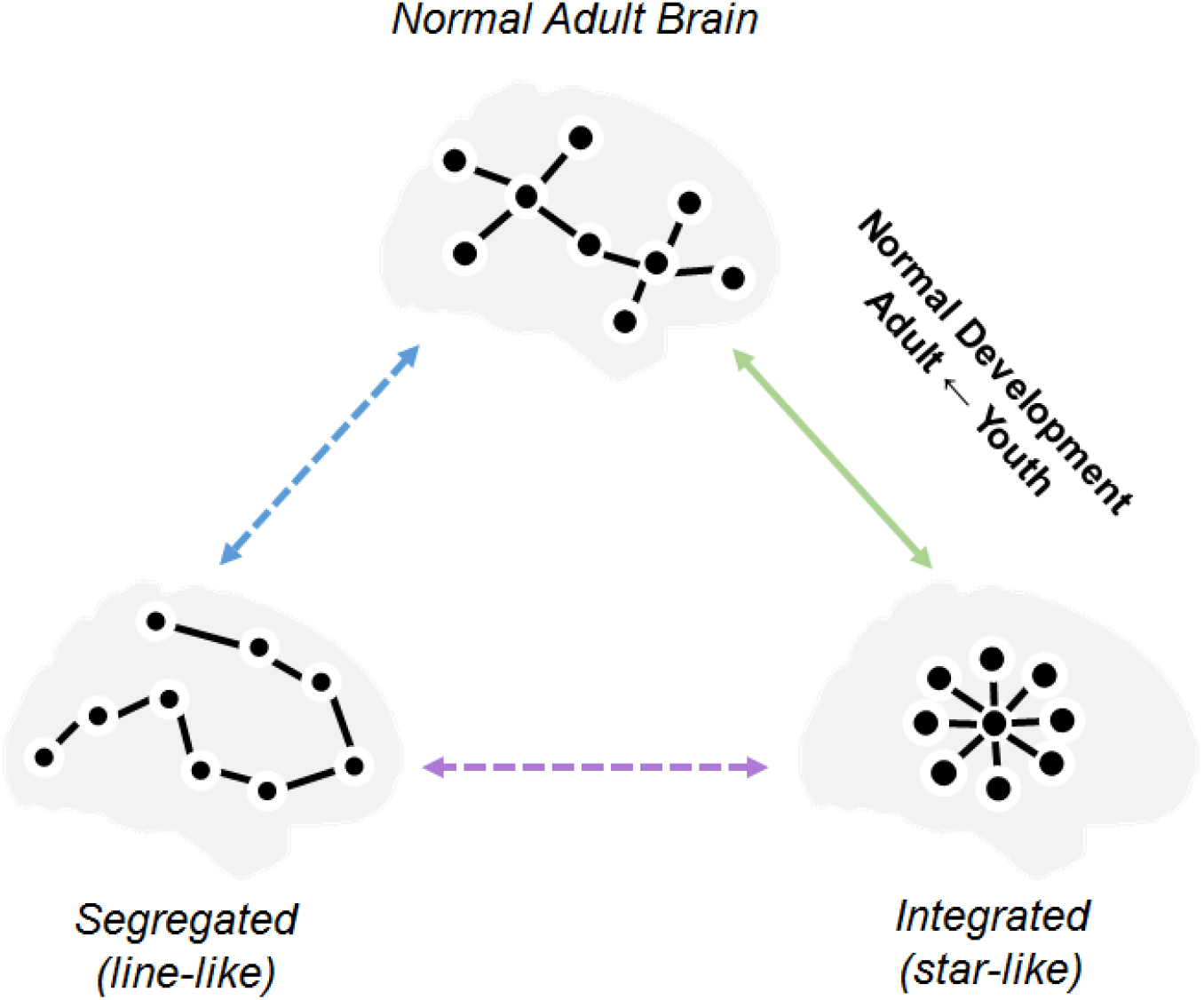
A heuristic minimum spanning tree (MST) model for the emergence of complex brain networks. This MST model space is based on the heuristic model of complex brain networks proposed by Stam and Van Straaten 2012. The model space consists of two extreme MSTs (representing network integration/segregation), an optimal MST for the normal adult brain, and three inter-connecting axes. Functional brain networks are proposed to develop from a star-like MST toward the optimal MST along the up-right axis, i.e., a balance between network integration and segregation. The solid line represents a developmental trajectory supported by this study, dashed lines represent trajectories that require future rigorous empirical support.

There are a few caveats worth mentioning in relation to the future application of this work. From a theoretical point of view, it is conceded that there are currently no simple mathematical models that fully characterise healthy brain networks, such as its hierarchical modularity, in order to fill the gap between the existing small-world and scale-free network models. Tree Hierarchy is a composite measure of network hierarchy, and thus is inherently correlated with other measures such as leaf number and maximum betweenness centrality (see methods for details). Therefore, discovery of new mathematical models will likely support a deeper understanding of network constraints on the developing brain (***Stam and van Straaten, 2012***). From a methodological point of view, although in the present study we took care of signal leakage in source space using the leakage-invariant PLI metric, and loops were discarded in the MST construction, the data may still have suffered to some extent from so-called secondary leakage (***Palva and Palva, 2012; Wang et al., 2018***). Therefore, future studies would also benefit from advanced methods such as implementing Lowdin Orthogonalisation (***Lowdin, 1950***) in MEG connectivity/network analyses to reduce those “ghost” connections (***Colclough et al., 2015***). Furthermore, for the warping procedure in children, we initially tested with age-specific paediatric templates as it was suspected that, in comparison to the adult template, the paediatric ones would produce a better approximation to the child’s brain anatomy due to better alignment in terms of skull thickness and brain morphology. However, adult and child templates produced very similar results in a previous study (***Cheyne et al., 2014***). Moreover, the AAL atlas was not available for these paediatric templates, hence using the anatomical labelling from the AAL (adult) atlas (***Tzourio-Mazoyer et al., 2002***) in paediatric surrogate structural MRI would still only provide an approximate labelling. Therefore, the surrogate procedure (using the adult template), as well as the subsequent analyses, were kept the same for all participants. Nevertheless, the use of age-specific template brain images and atlases together with surface-based registration in further studies would help to minimise registration errors due to the heterogeneity of brain anatomy in young children (***Fonov et al., 2011***). In addition, canonically-defined frequency bands may overlook some physiological mechanisms underlying the development of oscillatory neural networks. Estimating network properties from age-appropriate frequency bands is critical in future work (***Boersma et al., 2013***), for example by parameterization of neuronal power spectral densities on the basis of putative oscillatory components (***Haller et al., 2018***). Lastly, the developmental trajectory found in this cross-sectional study should be replicated in a large longitudinal sample.

In conclusion, a combination of an atlas-based beamformer in age-appropriate MEG data, leakage-insensitive PLI connectivity estimation, and unbiased MST network measures revealed that functional brain networks become more segregated during childhood. Increases in MST Diameter and decreases in Leaf Fraction indicate that functional networks develop into a more line-like (de-centralised) topology; increases in Degree Correlation and Eccentricity suggest that brain regions stay less central and become more locally specialised; decreases in Kappa and Tree Hierarchy emphasise that the network segregation during development balances the benefits of integration between distant brain regions against the risks of overload on central regions. Importantly, these topological network changes are most evident in the preschool years of childhood (i.e., the younger age group between 4-6 years in our data) and exhibit the same pattern for all ferquency bands (i.e., delta to low gamma). Our data resolves a long-standing debate in the field with respect to the normative brain development across spatial and temporal scales of investigation using MRI-based and electrophysiological measures. Finally, we propose a heuristic MST model for the emergence of complex brain networks, in which different patterns of network abnormality could be discerned depending upon their trajectories through this “network space”. Therefore, our study also represents the first attempt in providing a unifying network model for the development of functional brain networks in youth. We anticipate new data from both normative and abnormal developmental studies to be incorporated into this network space to enable us not only to understand new mechanisms for early brain development and resolve ambiguities in the field, but most importantly to translate brain network studies into solutions for clinical diagnosis and treatments.

## Methods and Materials

### Participants

Included participants were control participants who took part in a larger project on stuttering. The dataset consisted of MEG recordings collected from 28 children and 24 adults during 3-5 minutes of eyes-open resting-state. Due to excessive head movement, incidental system noise or signs of drowsiness, data from 4 children were excluded. The present analyses were therefore completed on a total of 48 participants: 24 children aged from 4 to 12 years, and 24 adults (*μ* = 40.6, *σ* = 17.4, 16 males). Children were further divided into two groups: a younger group with mean age centred at 5 years (5 Y.O., N = 10, *μ* = 5.4, *σ* = 1.1, 5 males) and an older group at 10 years (10 Y.O., N = 14, *μ* = 9.8, *σ* = 1.5, 12 males).

The experimental procedures were approved by the Human Participants Ethics Committee at Macquarie University. Written consent was obtained from the adult participants and from the parents/guardians of the children prior to the experiment. All participants were remunerated for their participation.

### Experimental Procedures

Upon arriving at the laboratory, participants were familiarised with the magnetically shielded room where they would be tested in a supine position. Prior to MEG measurements, five head position indicators (HPIs) were attached to a tightly fitting elastic cap. The 3D locations of the HPIs, fiducial landmarks (nasion, and left and right pre-auricular points) and the shape of each participant’s head were measured with a pen digitiser (Polhemus Fastrak, Colchester, VT, USA).

Children in the 5 Y.O. group were tested using the child-customized 125-channel whole-head gradiometer MEG system (Model PQ1064R-N2m, KIT, Kanazawa, Japan), and all other participants were tested using the 160-channel whole-head gradiometer MEG system (Model PQ1160RN2, KIT, Kanazawa, Japan). The gradiometers of both systems have a 50 mm baseline and 15.5 mm diameter coils, and are positioned in a glass fibre reinforced plastic cryostat for measurement of the normal component of the magnetic field from the human brain (***Kado et al., 1999***). In both systems, neighbouring channels are 38 mm apart and 20 mm from the outer dewar surface. The 125-channel dewar was designed to fit a maximum head circumference of 53.4 cm, accommodating more than 90% of heads of 5-year olds ***Johnson et al., 2010***). Both systems were situated within the same magnetically shielded room, and therefore have comparable environmental noise level.

During MEG data acquisition, participants were asked to remain relaxed, awake and with their eyes fixed on a white cross at the centre of a black 36 cm (width) x 24 cm (length) rectangular image with 4×4 degrees of visual angle. The visual presentation was done by video projectors situated outside the magnetically shielded room (child MEG projector: Sharp Notevision Model PG10S, Osaka, Japan; Adult MEG projector: InFocus Model IN5108, Portland, USA). Drowsiness was monitored online through a video-camera so that any affected data would be removed from further analysis. For child participants, an experienced researcher sat with them during the whole session to make sure they were comfortable.

### MEG Data Pre-processing

MEG data were acquired at a sampling frequency of 1000 Hz and with an online bandpass of 0.03-200 Hz. Head positions were measured at the beginning and end of the acquisition session; a movement tolerance of 5 mm and 10 mm was used in adults and children, respectively.

The Yokogawa/KIT MEG data were firstly converted to a CTF data format using BrainWave toolbox developed at the Hospital for Sick Children in Canada (http://cheynelab.utoronto.ca, version 3.3beta, see Cheyne et al., 2014 for details). Then, the CTF compatible MEG data were imported into and processed using DataEditor in the CTF MEG5 software (VSM MedTech Systems Inc., Coquitlam BC, Canada; Version 5.0.2). The continuous raw MEG data were firstly filtered from 0.5 to 100 Hz using bi-directional IIR Butterworth filters with DC removal and segmented into epochs of 4096 samples (= 4.096 seconds). Epochs that contained physiological (e.g., muscle noise) or environmental artefacts were rejected by visual inspection. The cleaned datasets consisted on average of 23.8 (*σ* = 3.02) epochs for the children and 40 epochs (*σ* = 0.02) for the adults.

### Head Modelling and Surrogate MRIs

For the head model construction, obtaining individual structural MRI scans of children - especially of those aged below 6 years - was impractical. A “surrogate” MRI approach was therefore used here to warp the adult Montreal Neurological Institute (MNI) template T1 structural brain image to each participant’s digitized head shape with an iterative closest point algorithm implemented in BrainWave (see ***Cheyne et al. 2014*** for details). MEG data was co-registered with the warped “surrogate” MRI using the digitised fiducial points. The outline of the scalp from this co-registered “surrogate” MRI was extracted using the MRIViewer in the CTF MEG5 software (VSM MedTech Systems Inc., Coquitlam BC, Canada; Version 5.0.2) and then used to fit a multisphere volume conductor model (***Huang et al., 1999***), which was subsequently used for the beamformer analysis described below.

### Beamforming

An atlas-based beamforming approach (***Hillebrand et al., 2012***) was adopted to project sensor level MEG data to source space. The co-registered surrogate MRIs were normalised to the standard MNI (T1) template, using the SEG toolbox (***Weiskopf et al., 2011***) in SPM8. The automated anatomical labelling (AAL) atlas (***Tzourio-Mazoyer et al., 2002***) was used to label the voxels in a participant’s normalised co-registered surrogate MRI, following which the centroid for each AAL regions of interest (80 ROIs; 78 cortical and bilateral hippocampal) was inversely transformed to native space (***Hillebrand et al., 2016***).

For each centroid, beamformer weights were computed using Synthetic Aperture Magnetometry (SAM, ***Robinson 1999***. This beamformer selectively weights the contribution from each MEG sensor to a voxel’s activity based on the broad-band (0.5-48 Hz) data covariance matrix, which was computed from (1) all selected time-series, (2) the forward solution (lead field) for a dipolar source with optimum orientation at that location, and (3) a unity noise covariance that was scaled by the smallest singular value in a decomposition of the data covariance matrix. The broad-band MEG data were subsequently projected through the normalised beamformer weights***Cheyne et al. (2007)***.

From the resulting time-series, the first 15 artifact-free epochs, containing 4096 samples (= 4.096 seconds), were selected for further analyses of functional connectivity and network topology. These selected epochs were then band-pass filtered, using an offline discrete Fast Fourier Transform filter without phase distortion, as implemented in the BrainWave toolbox developed at VU University Medical Centre (C.J. Stam; http://home.kpn.nl/stam7883/brainwave.html, version 0.9.152.4.1), into five canonical MEG frequency bands (delta: 0.5–4 Hz, theta: 4–8 Hz, alpha: 8–13 Hz, beta: 13–30 Hz, and low gamma: 30-48 Hz). Subsequently, the instantaneous phase for each time-series was determined by taking the argument of the analytic signal as computed using the Hilbert transform (Marple, 1999).

### Connectivity Analysis

Pair-wise frequency band-specific functional connectivity between the 80 ROIs was estimated using the phase lag index (PLI) for each of the 15 artifact-free epochs (= 4.096 seconds). PLI reflects the consistency by which one signal is phase leading or lagging with respect to another signal (***Stam et al., 2007***), which can be expressed as:

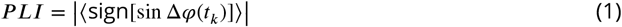

where Δ*φ* refers to the instantaneous phase difference between two time-series, *t_k_* are discrete time steps calculated over all ***K*** = 1 … *N*, *sign* refers to the signum function, <> and || denote the mean and absolute value, respectively. Specifically, PLI quantifies phase synchronisation as a measure of the asymmetry in the distribution of instantaneous phase differences between two time-series (in our case the beamformer reconstructed time-series for two ROIs). The value of PLI ranges from zero (random phase differences/no functional connectivity or only zero-lag/mod π) and one (perfect non-zero-lag synchrony). Because the effects of volume conduction/field spread/signal leakage give zero-lag (mod π) phase differences, PLI is insensitive to these effects at the cost of being blind to true zero-lag interactions. For each frequency band and each epoch, the 80 x 80 connectivity matrix of pairwise PLI values was computed. ROI-PLI was computed as the average PLI between a node and all other nodes, and whole-brain PLI was calculated as the average across all nodal PLI values.

### Minimum Spanning Tree Analysis

For each epoch and participant separately, the minimum spanning tree (MST) sub-graph was constructed using the PLI connectivity matrix. The MST is constructed by connecting all *n* nodes in such a way that the cost (the sum of all link weights) is minimised without forming cycles. For the computation of the MST, 1/PLI is used as the link weights since we are interested in the strongest connections in the network. MSTs were constructed in BrainWave by applying Kruskal’s algorithm (***Kruskal, 1956***), which starts with an unconnected network, adds the link with lowest weight, then adds the link with next lowest weight (if this does not create a loop), until all nodes are connected, thereby forming a tree consisting of *m = n* − 1 links.

Two extreme tree topologies exist: (1) a line-like tree (A in *Figure 1*) where all nodes are connected to two other nodes with the exception of the two so-called “leaf-nodes” at either end that have only one link, and (2) a star-like tree (C in ***Figure 1***) where all leaves are connected to one central node. There are many different tree types between these two extremes (e.g., B in ***Figure 1***). The tree topology can be characterised with various measures (***Boersma et al., 2013***).

Global MST network measures are informative about the functional integration and segregation of the entire network. Five different global MST measures were used here: (1) the “Leaf Fraction” is computed as the number of leaf nodes, divided by the total number of nodes; (2) the “Diameter” is the longest shortest path between any two nodes, where the shortest path is defined as the path with smallest number of links between two nodes; (3) the “Tree Hierarchy” was introduced (***Boersma et al., 2013***) to describe a balance between a small diameter without overloading central nodes in the tree (***Figure 1***). It is defined as 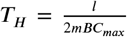, where *l* is the leaf number and ***BC_max_*** represents the maximal betweenness centrality in the tree. In a line-like tree, *l* = 2 and with m approaching infinity, *T_H_* approaches 0; and in a star-like tree, *l* ≈ *m*, so *T_H_* approaches 0.5; for *l* between these two extremes, *T_H_* can have higher values (with an upper bound of 1); (4) the “Degree Correlation” is an index of whether the degree of a node is correlated with the degree of its neighbouring nodes (***Van Mieghem et al., 2010***); (5) “Kappa” (also called degree divergence; ***Barrat et al. 2008***) measures the broadness of the degree distribution, and is high in graphs with a scale-free degree distribution, and low in graphs with a degree distribution that approaches the normal distribution. Kappa also relates to network robustness: high kappa reflects high resilience against random damage in networks.

Nodal MST network measures capture the importance of a node within the network. Three different nodal measures for centrality (“hubness”) were used: (1) the “Degree” is the number of connections of a node to its neighbouring nodes; (2) the “Betweenness Centrality” is the fraction of the shortest paths that pass through a node; (3) the “Eccentricity” of a node is the longest shortest path between a node and any other node, and is low if the node is central in the graph (***Bullmore and Sporns, 2012***).

### Statistical Analysis

Statistical analyses were performed using permutation testing as implemented in the Resampling Statistical Toolkit for Matlab 2016a. We used 50,000 permutations of group membership to empirically approximate the distribution for the null hypothesis (i.e., no difference between groups) for each contrast. For each permutation, the F/t values were derived for a contrast of interest, and any F/t values for the original data that exceeded the significance threshold for the F/t distribution were deemed reliable. Furthermore, *p* values were corrected for multiple comparisons at the threshold of 0.05 using the false discovery rate (FDR, ***Benjamini and Hochberg 1995***).

For each frequency band and each participant separately, whole-brain PLI were averaged over the 15 epochs per participant. The ROI-PLI values, global and nodal MST measures were averaged over 15 epochs, yielding 80 ROI-PLI, 5 global MST, and 3 x 80 (= nodal MST measures x ROIs) values per participant for each frequency band, respectively.

Permutation tests were initially performed, for each frequency band separately, between adults and children (as a whole group), for the whole-brain PLI and the global MST measures (FDR corrected for the number of global measures (5)); if the whole-brain PLI or the global MST measures were significantly different in a specific frequency band, then the ROI-PLI and the nodal MST measures were compared (FDR corrected for three nodal measures x 80 ROIs). Second level permutation tests were performed in pairwise groups (10 Y.O. versus 5 Y.O., adults versus 5 Y.O., adults versus 10 Y.O.) for the whole-brain PLI or the global MST measures if adults and children (as a whole group) showed significant differences for these measures in any specific frequency band, and for the ROI-PLI or the nodal MST measures if these measures were significantly different in any specific frequency band between adults and children (as a whole group).

## Acknowledgments

We thank all participants for their participation. We also thank Douglas Cheyne and Cecilia Jobst for their assistance in the preliminary data analysis. Finally, we acknowledge the collaboration of Kanazawa Institute of Technology in establishing the KIT-Macquarie MEG laboratory.

## Additional information

### Author contributions

Wei He: conceptualisation, funding acquisition, data curation, formal analysis, validation, visualisation, methodology, writing (original draft, reviewing and editing); Paul F Sowman: conceptualisation, funding acquisition, data collection, writing (reviewing and editing); Jon Brock: conceptualisation, funding acquisition, writing (reviewing and editing); Andrew C. Etchell: project management, data collection and curation; Cornelis J. Stam: conceptualisation, resources, methodology, writing (reviewing and editing); Arjan Hillebrand: conceptualisation, resources, formal analysis, validation, visualisation, methodology, supervision, writing (original draft, reviewing and editing).

## Appendix 1

**Appendix 1 Table 1.**
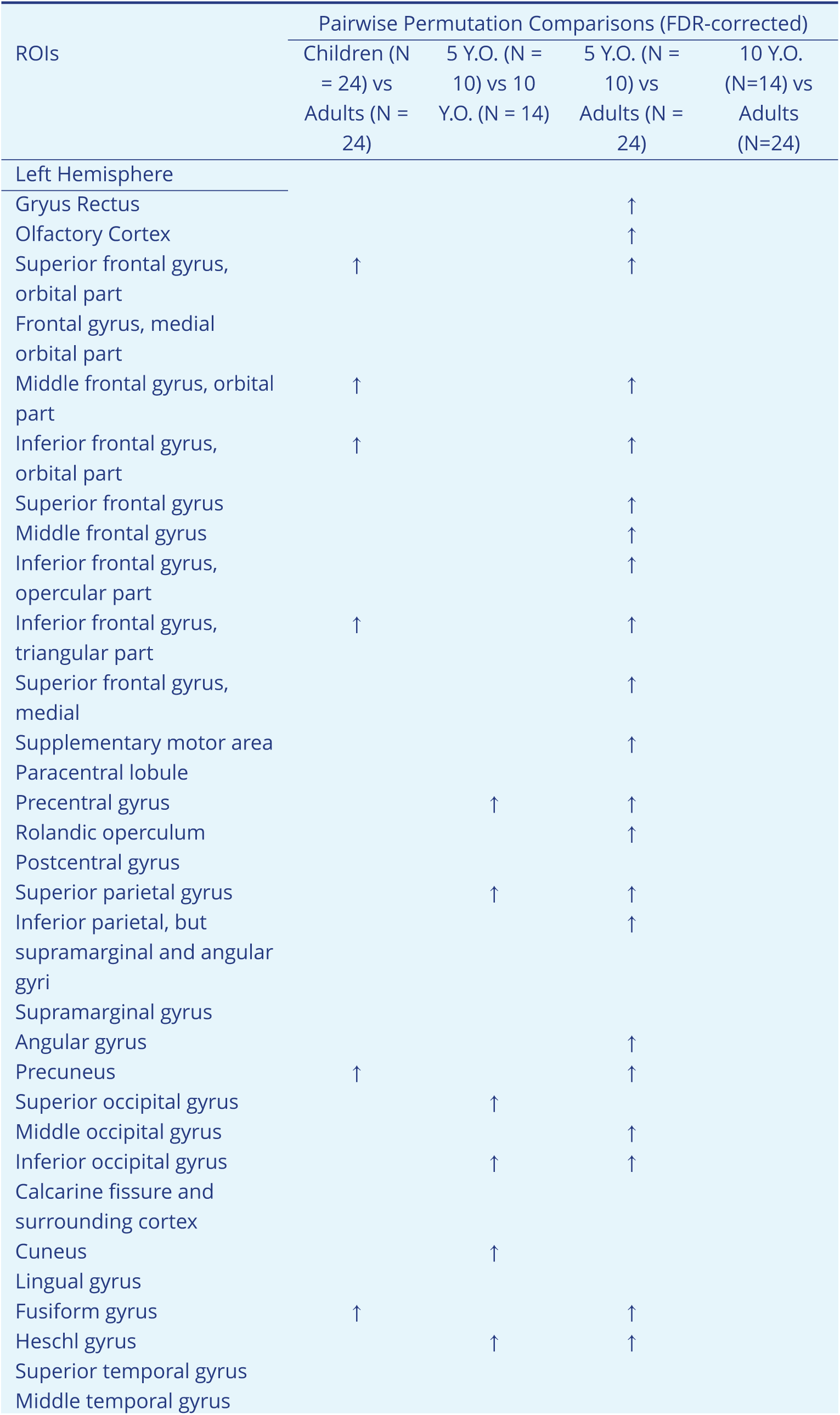
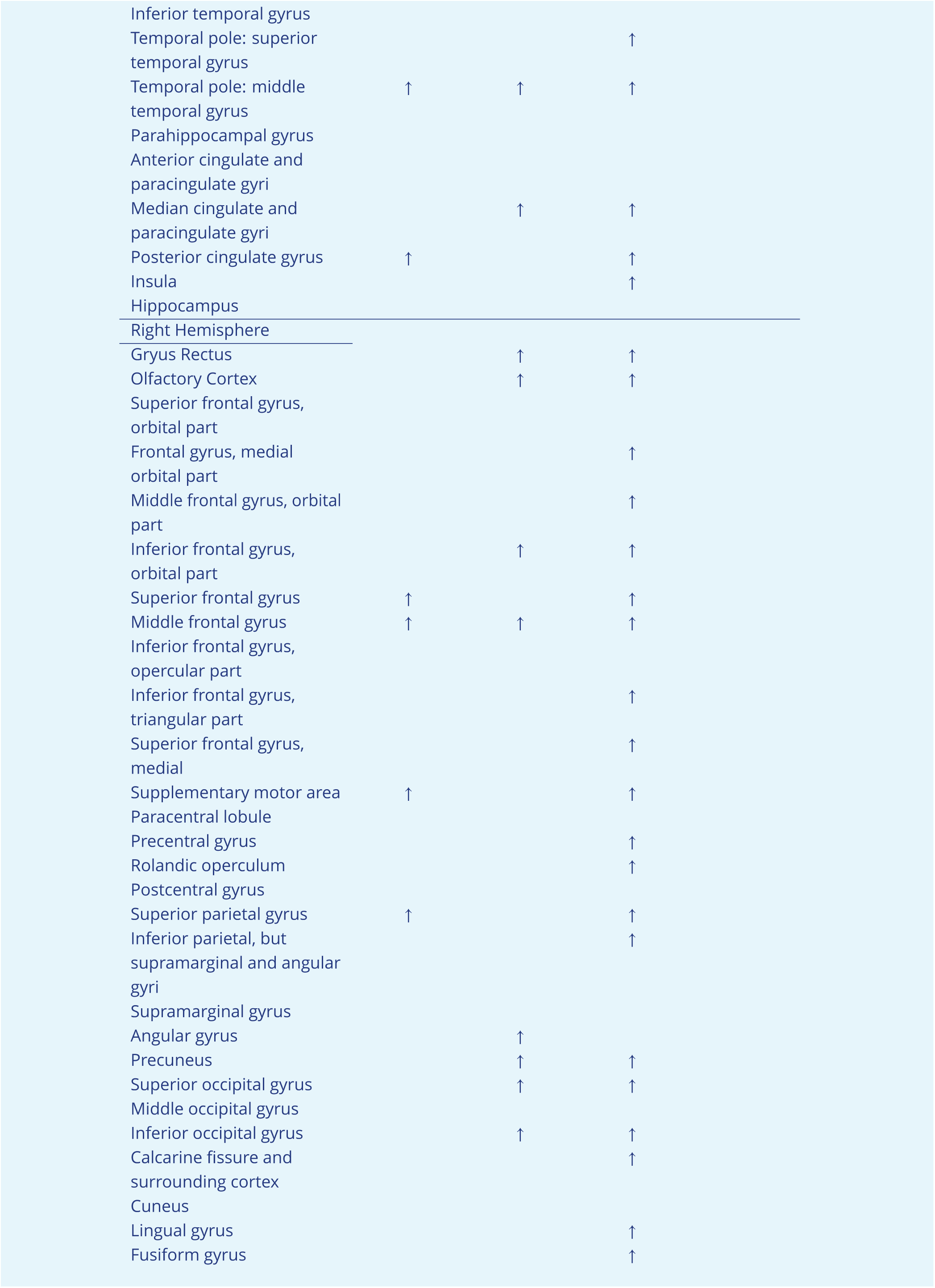
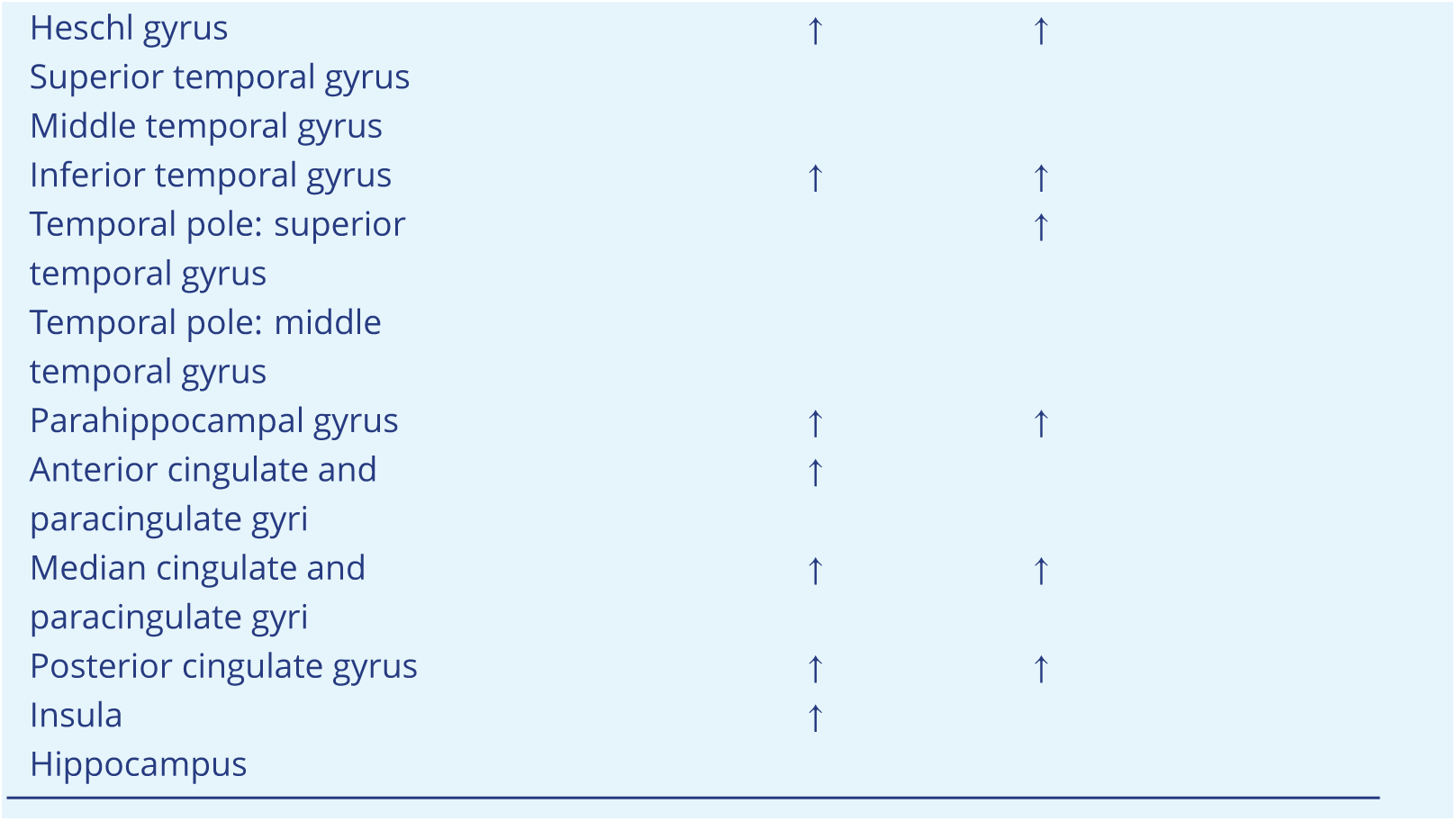
Regions of interest (ROIs) that manifest significant Eccentricity differences between groups in the delta band.

**Appendix 1 Table 2.**
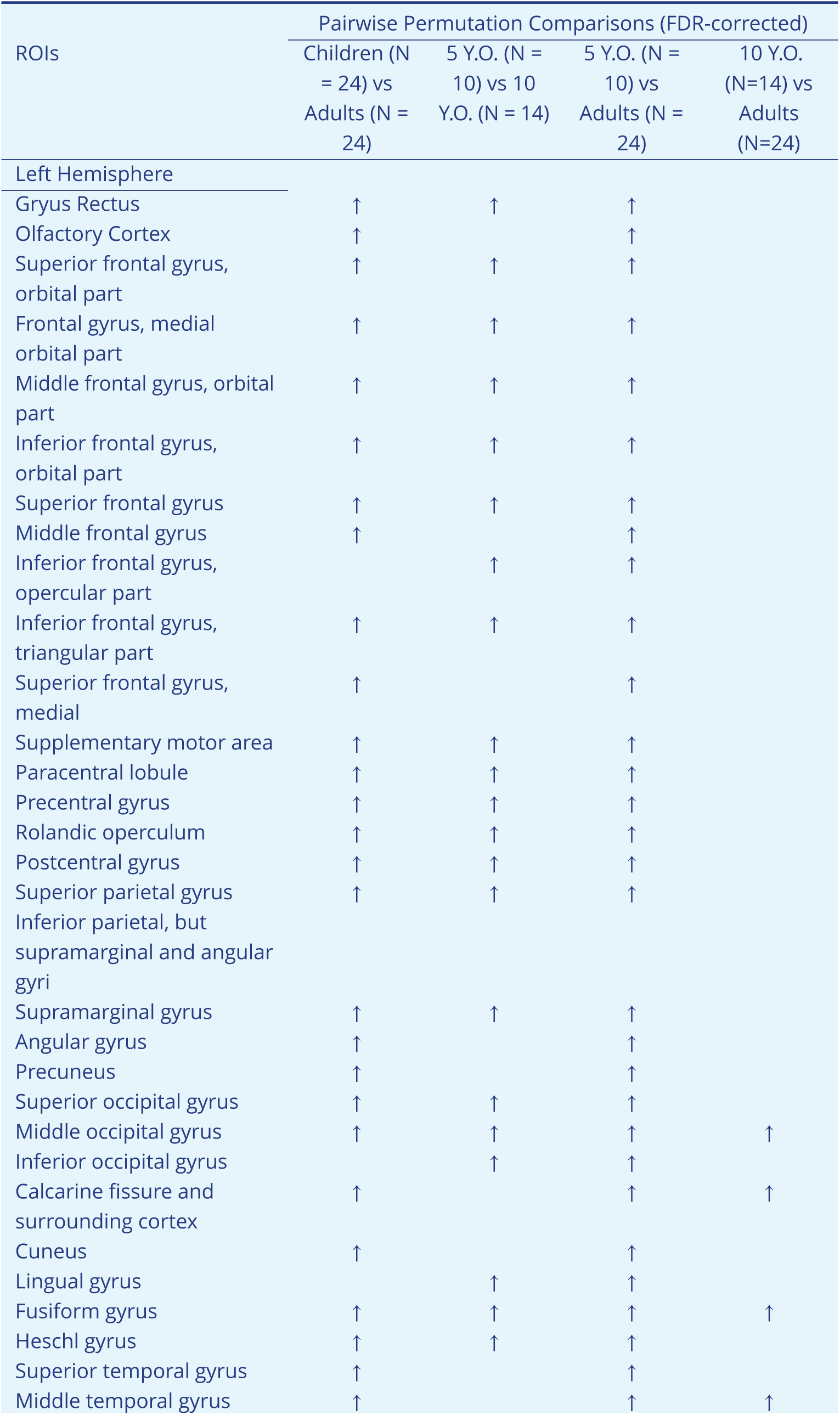
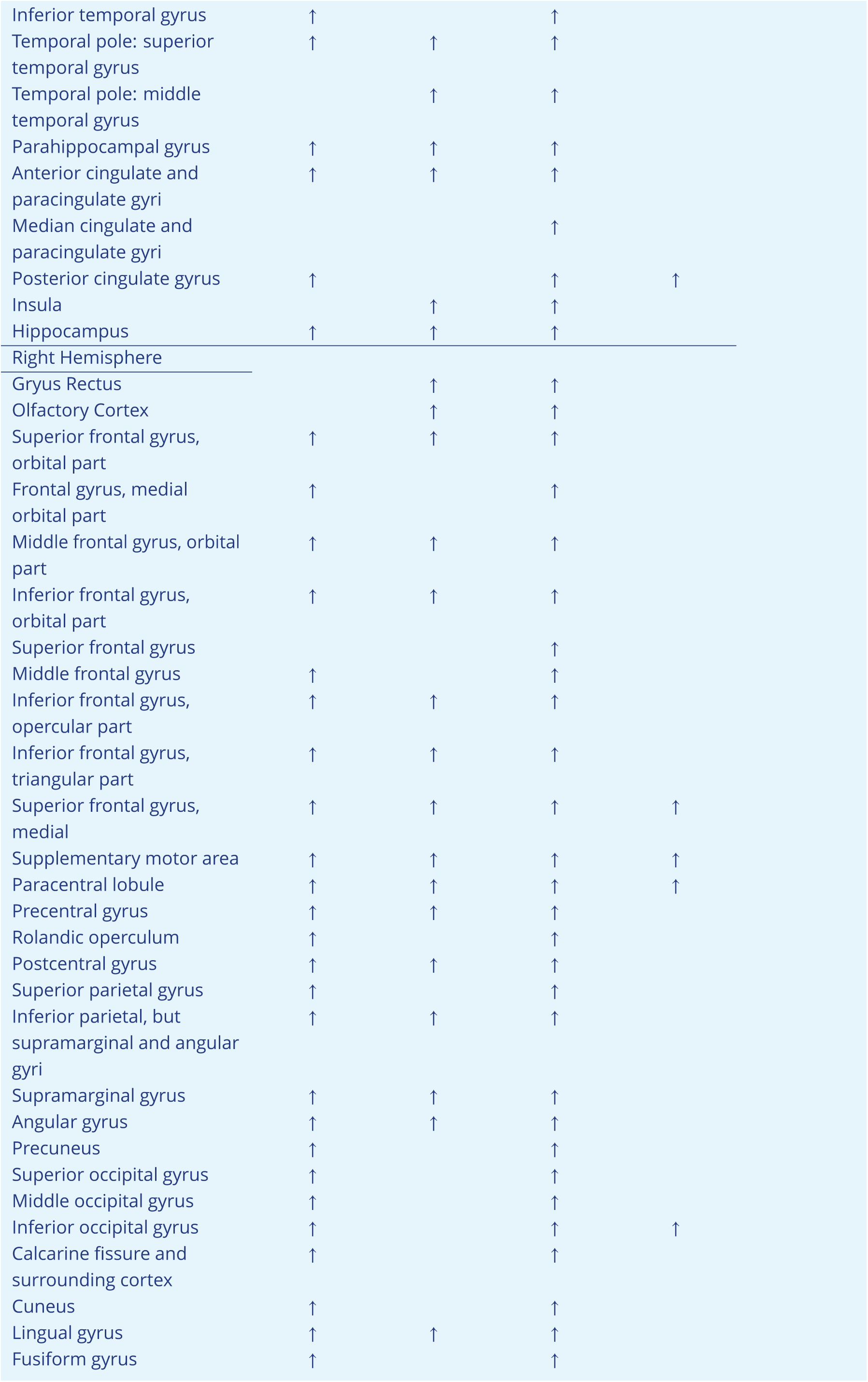
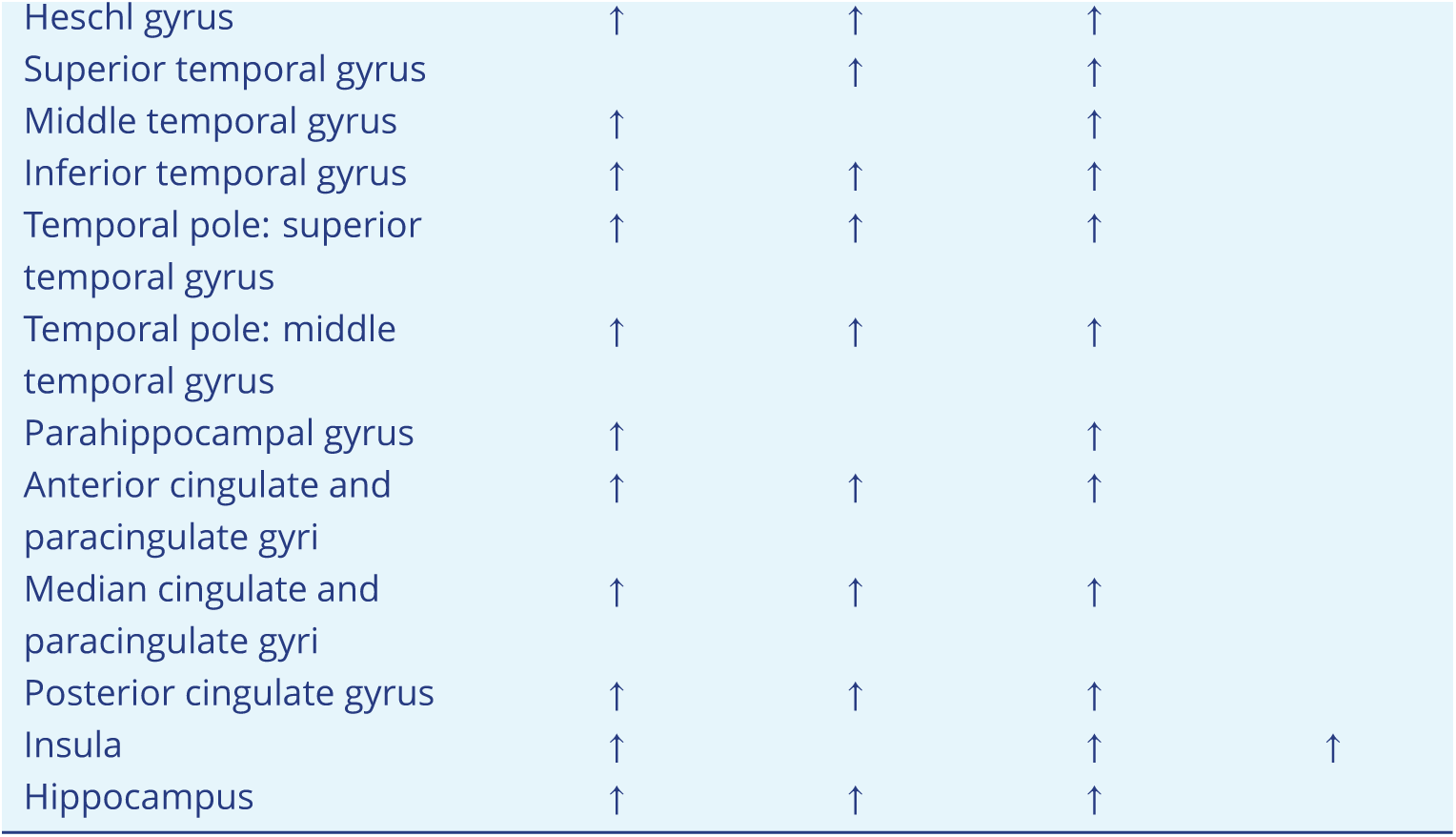
Regions of interest (ROIs) that manifest significant Eccentricity differences between groups in the theta band.

**Appendix 1 Table 3.**
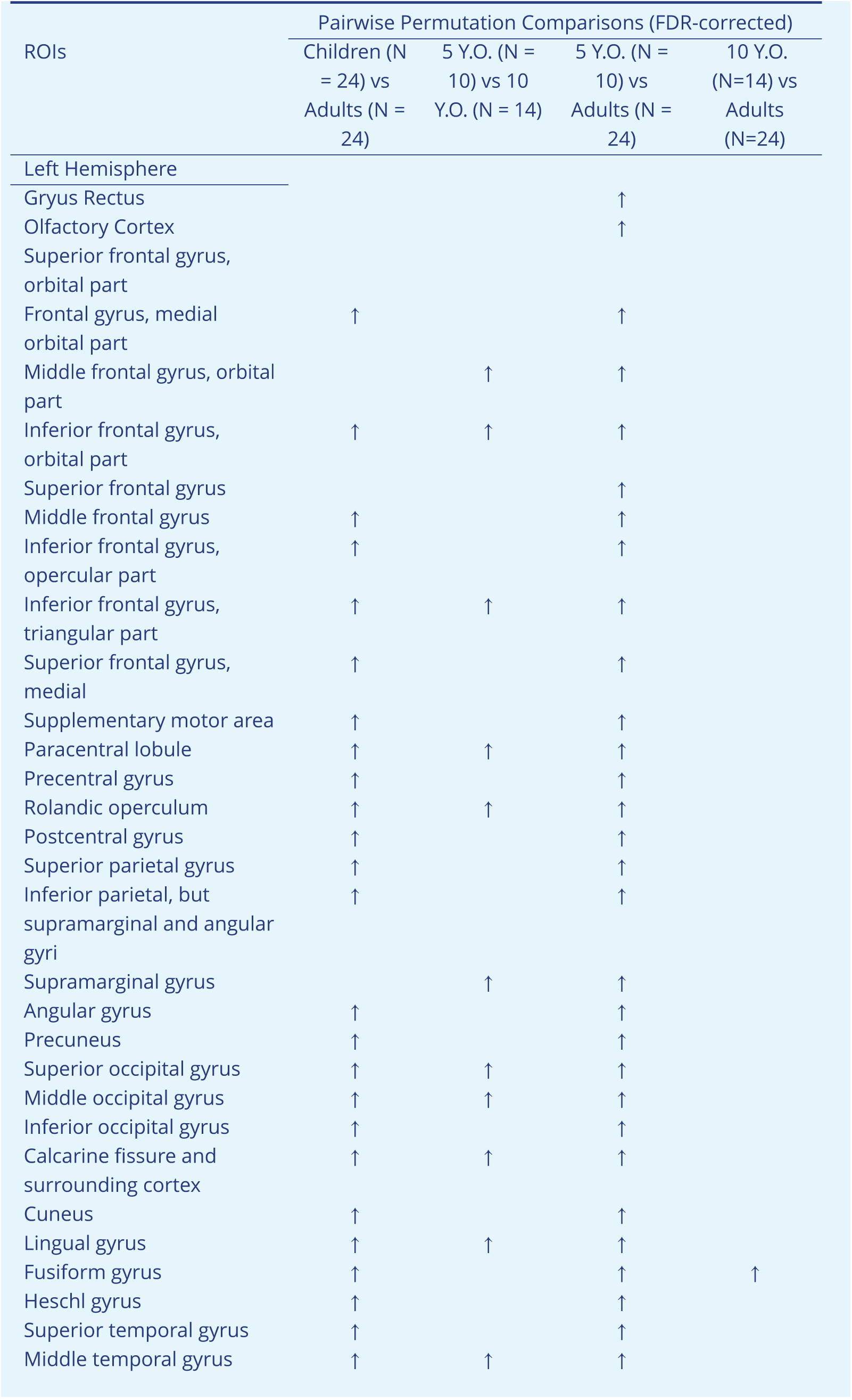
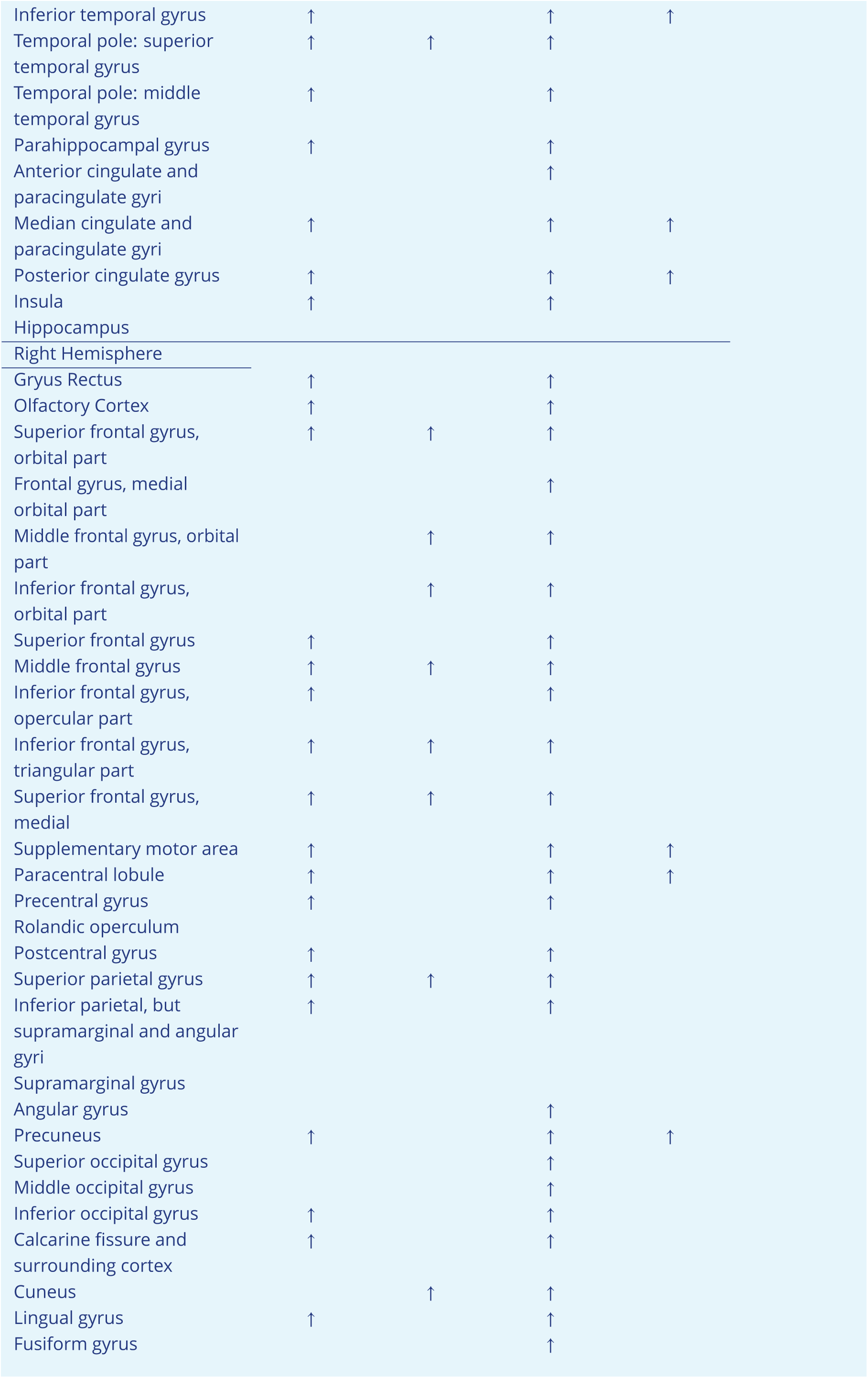
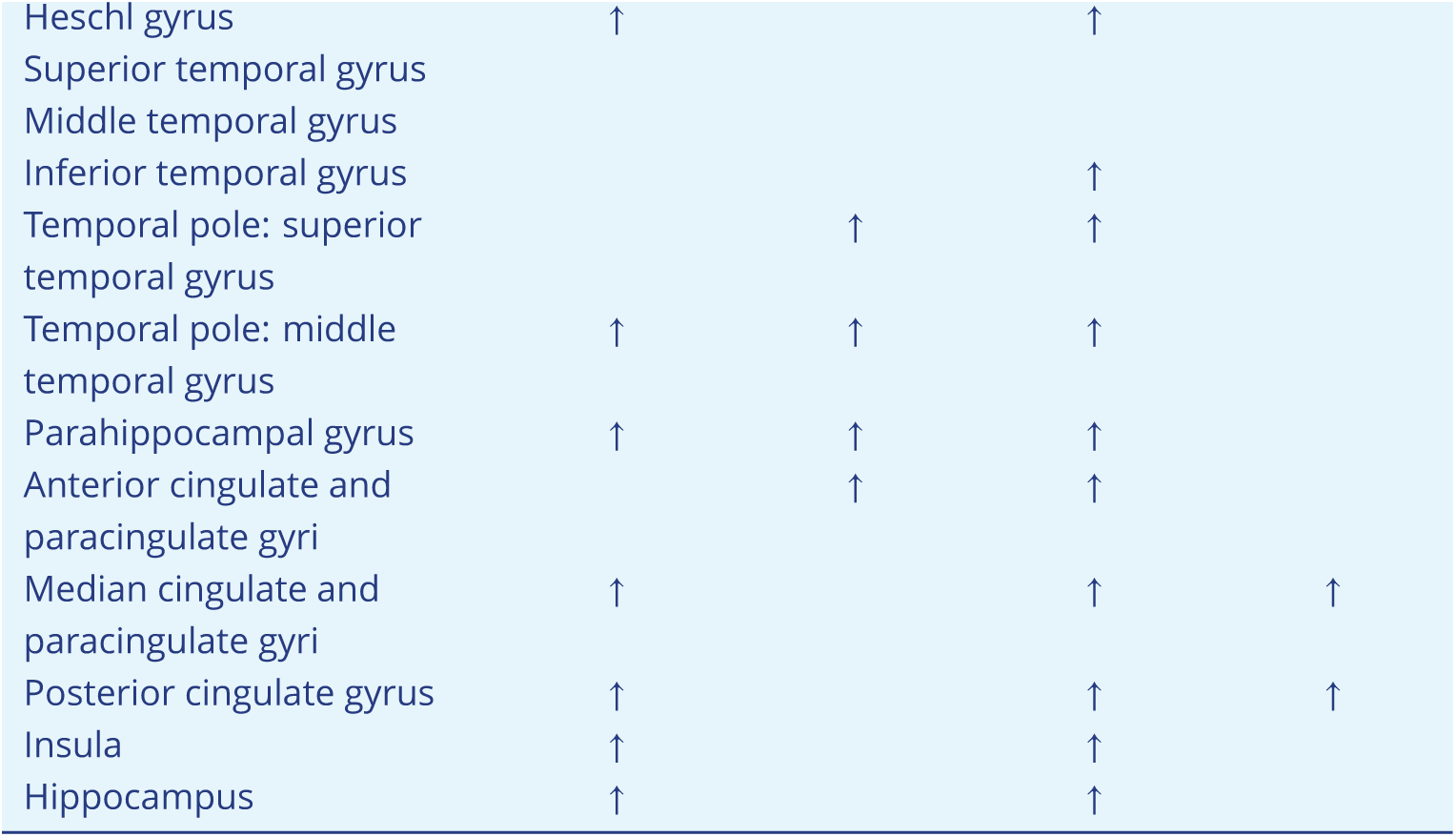
Regions of interest (ROIs) that manifest significant Eccentricity differences between groups in the alpha band.

**Appendix 1 Table 4.**
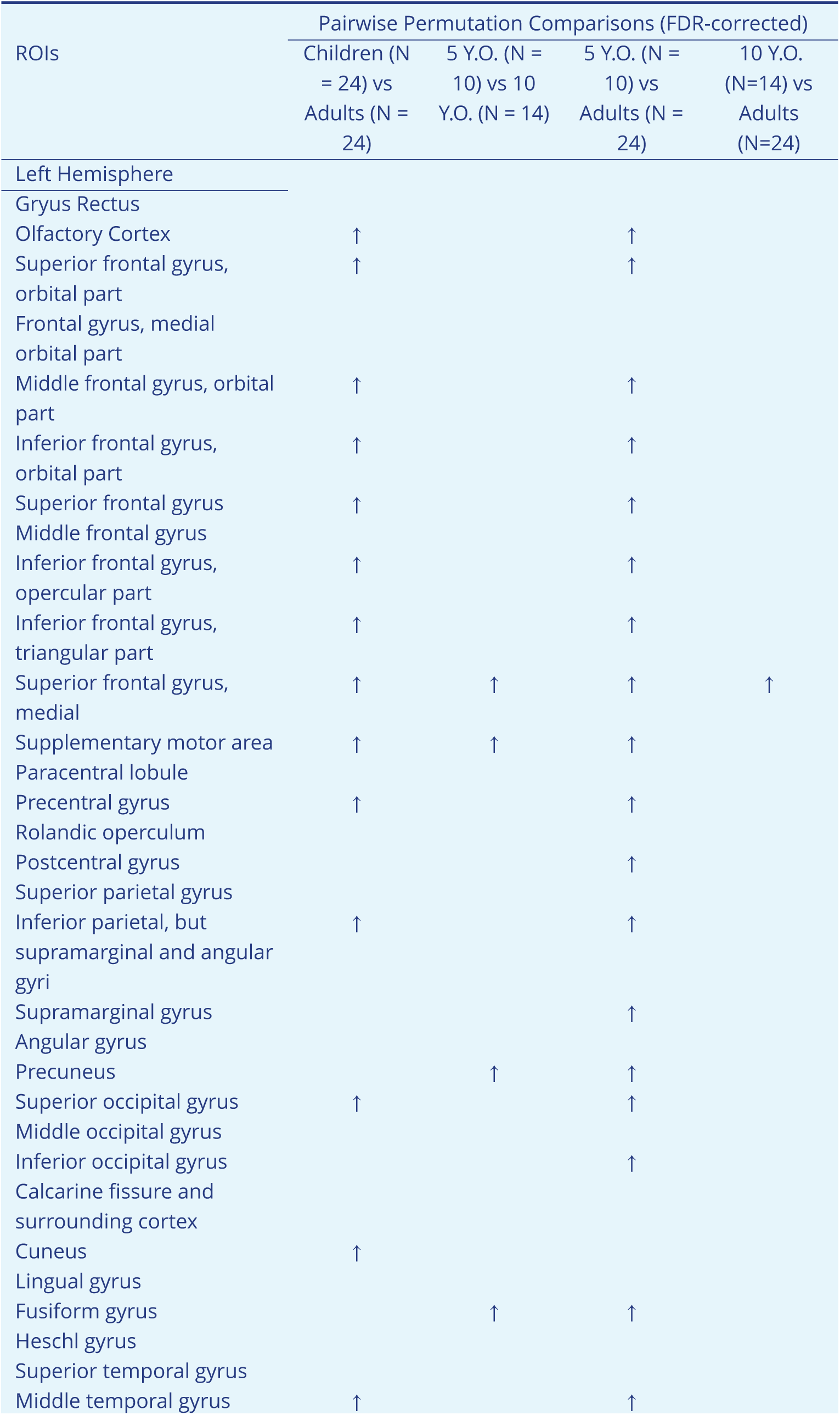
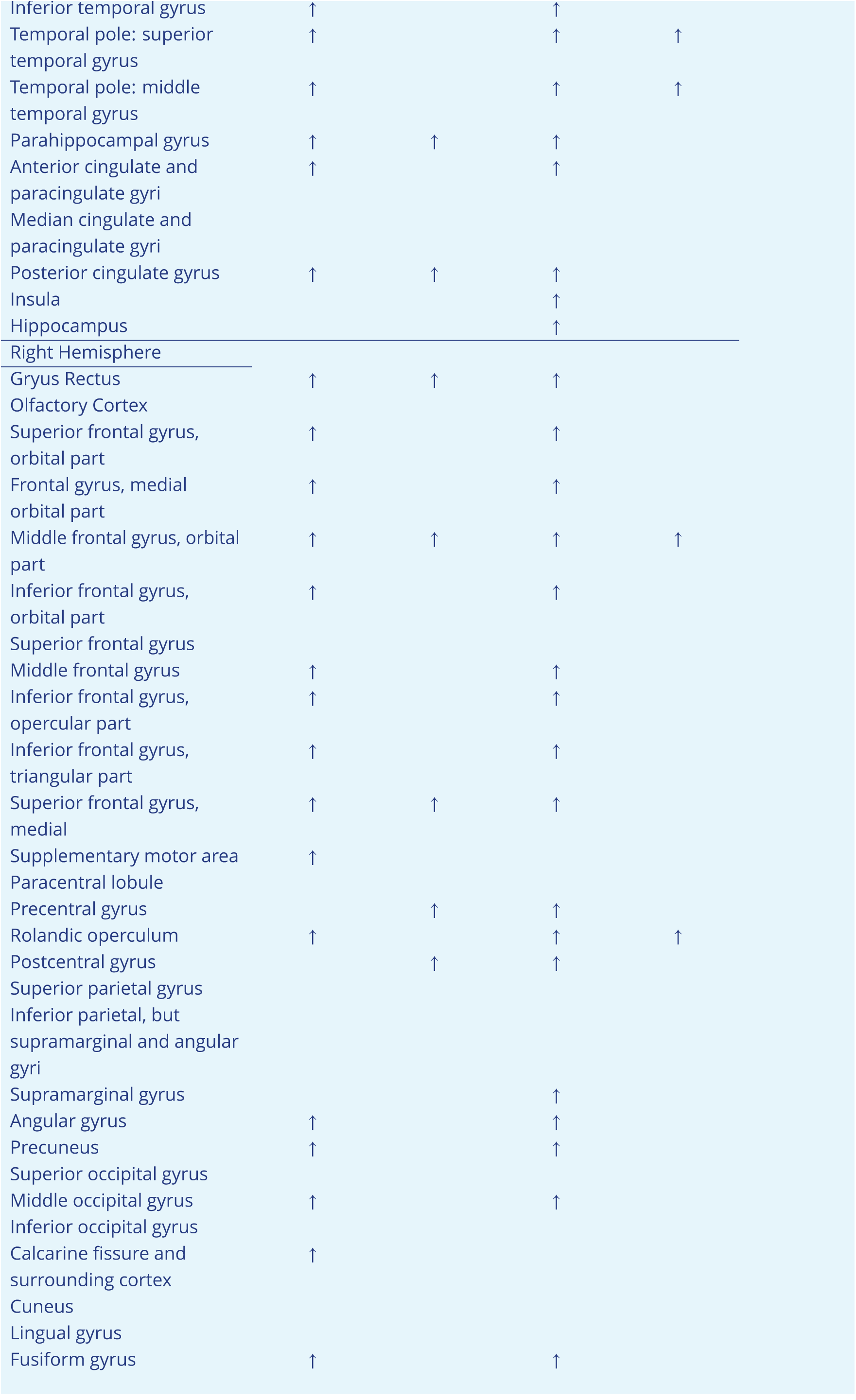
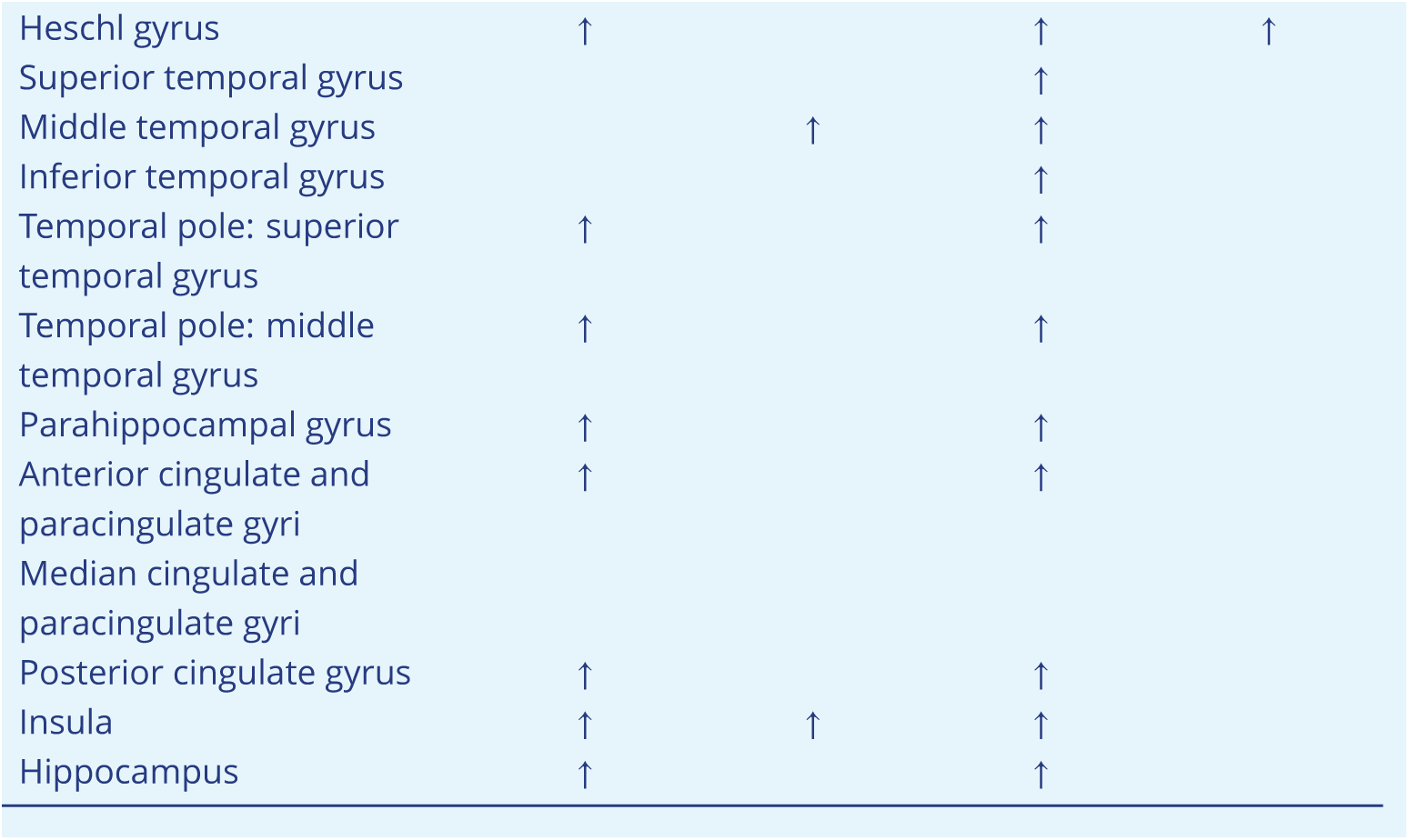
Regions of interest (ROIs) that manifest significant Eccentricity differences between groups in the beta band.

| ROIs | Pairwise Permutation Comparisons (FDR-corrected) |  |  |  |
| --- | --- | --- | --- | --- |
|  | Children (N = 24) vs Adults (N = 24) | 5 Y.O. (N = 10) vs 10 Y.O. (N = 14) | 5 Y.O. (N = 10) vs Adults (N = 24) | 10 Y.O. (N=14) vs Adults (N=24) |
| Left Hemisphere |  |  |  |  |
| Gyrus Rectus |  |  |  |  |
| Olfactory Cortex | ↑ |  | ↑ |  |
| Superior frontal gyrus, orbital part | ↑ |  | ↑ |  |
| Frontal gyrus, medial orbital part |  |  |  |  |
| Middle frontal gyrus, orbital part | ↑ |  | ↑ |  |
| Inferior frontal gyrus, orbital part | ↑ |  | ↑ |  |
| Superior frontal gyrus | ↑ |  | ↑ |  |
| Middle frontal gyrus |  |  |  |  |
| Inferior frontal gyrus, opercular part | ↑ |  | ↑ |  |
| Inferior frontal gyrus, triangular part | ↑ |  | ↑ |  |
| Superior frontal gyrus, medial | ↑ | ↑ | ↑ | ↑ |
| Supplementary motor area | ↑ | ↑ | ↑ |  |
| Paracentral lobule |  |  |  |  |
| Precentral gyrus | ↑ |  | ↑ |  |
| Rolandic operculum |  |  |  |  |
| Postcentral gyrus |  |  | ↑ |  |
| Superior parietal gyrus |  |  |  |  |
| Inferior parietal, but supramarginal and angular gyri | ↑ |  | ↑ |  |
| Supramarginal gyrus |  |  | ↑ |  |
| Angular gyrus |  |  |  |  |
| Precuneus |  | ↑ | ↑ |  |
| Superior occipital gyrus | ↑ |  | ↑ |  |
| Middle occipital gyrus |  |  |  |  |
| Inferior occipital gyrus |  |  | ↑ |  |
| Calcarine fissure and surrounding cortex |  |  |  |  |
| Cuneus | ↑ |  |  |  |
| Lingual gyrus |  |  |  |  |
| Fusiform gyrus |  | ↑ | ↑ |  |
| Heschl gyrus |  |  |  |  |
| Superior temporal gyrus |  |  |  |  |
| Middle temporal gyrus | ↑ |  | ↑ |  |
| Inferior temporal gyrus | ↑ |  | ↑ |  |
| Temporal pole: superior temporal gyrus | ↑ |  | ↑ | ↑ |
| Temporal pole: middle temporal gyrus | ↑ |  | ↑ | ↑ |
| Parahippocampal gyrus | ↑ | ↑ | ↑ |  |
| Anterior cingulate and paracingulate gyri | ↑ |  | ↑ |  |
| Median cingulate and paracingulate gyri |  |  |  |  |
| Posterior cingulate gyrus | ↑ | ↑ | ↑ |  |
| Insula |  |  | ↑ |  |
| Hippocampus |  |  | ↑ |  |
| Right Hemisphere |  |  |  |  |
| Gyrus Rectus | ↑ | ↑ | ↑ |  |
| Olfactory Cortex |  |  |  |  |
| Superior frontal gyrus, orbital part | ↑ |  | ↑ |  |
| Frontal gyrus, medial orbital part | ↑ |  | ↑ |  |
| Middle frontal gyrus, orbital part | ↑ | ↑ | ↑ | ↑ |
| Inferior frontal gyrus, orbital part | ↑ |  | ↑ |  |
| Superior frontal gyrus |  |  |  |  |
| Middle frontal gyrus | ↑ |  | ↑ |  |
| Inferior frontal gyrus, opercular part | ↑ |  | ↑ |  |
| Inferior frontal gyrus, triangular part | ↑ |  | ↑ |  |
| Superior frontal gyrus, medial | ↑ | ↑ | ↑ |  |
| Supplementary motor area | ↑ |  |  |  |
| Paracentral lobule |  |  |  |  |
| Precentral gyrus |  | ↑ | ↑ |  |
| Rolandic operculum | ↑ |  | ↑ | ↑ |
| Postcentral gyrus |  | ↑ | ↑ |  |
| Superior parietal gyrus |  |  |  |  |
| Inferior parietal, but supramarginal and angular gyri |  |  |  |  |
| Supramarginal gyrus |  |  | ↑ |  |
| Angular gyrus | ↑ |  | ↑ |  |
| Precuneus | ↑ |  | ↑ |  |
| Superior occipital gyrus |  |  |  |  |
| Middle occipital gyrus | ↑ |  | ↑ |  |
| Inferior occipital gyrus |  |  |  |  |
| Calcarine fissure and surrounding cortex | ↑ |  |  |  |
| Cuneus |  |  |  |  |
| Lingual gyrus |  |  |  |  |
| Fusiform gyrus | ↑ |  | ↑ |  |
| Heschl gyrus | ↑ |  | ↑ | ↑ |
| Superior temporal gyrus |  |  | ↑ |  |
| Middle temporal gyrus |  | ↑ | ↑ |  |
| Inferior temporal gyrus |  |  | ↑ |  |
| Temporal pole: superior temporal gyrus | ↑ |  | ↑ |  |
| Temporal pole: middle temporal gyrus | ↑ |  | ↑ |  |
| Parahippocampal gyrus | ↑ |  | ↑ |  |
| Anterior cingulate and paracingulate gyri | ↑ |  | ↑ |  |
| Median cingulate and paracingulate gyri |  |  |  |  |
| Posterior cingulate gyrus | ↑ |  | ↑ |  |
| Insula | ↑ | ↑ | ↑ |  |
| Hippocampus | ↑ |  | ↑ |  |

**Appendix 1 Table 5.**
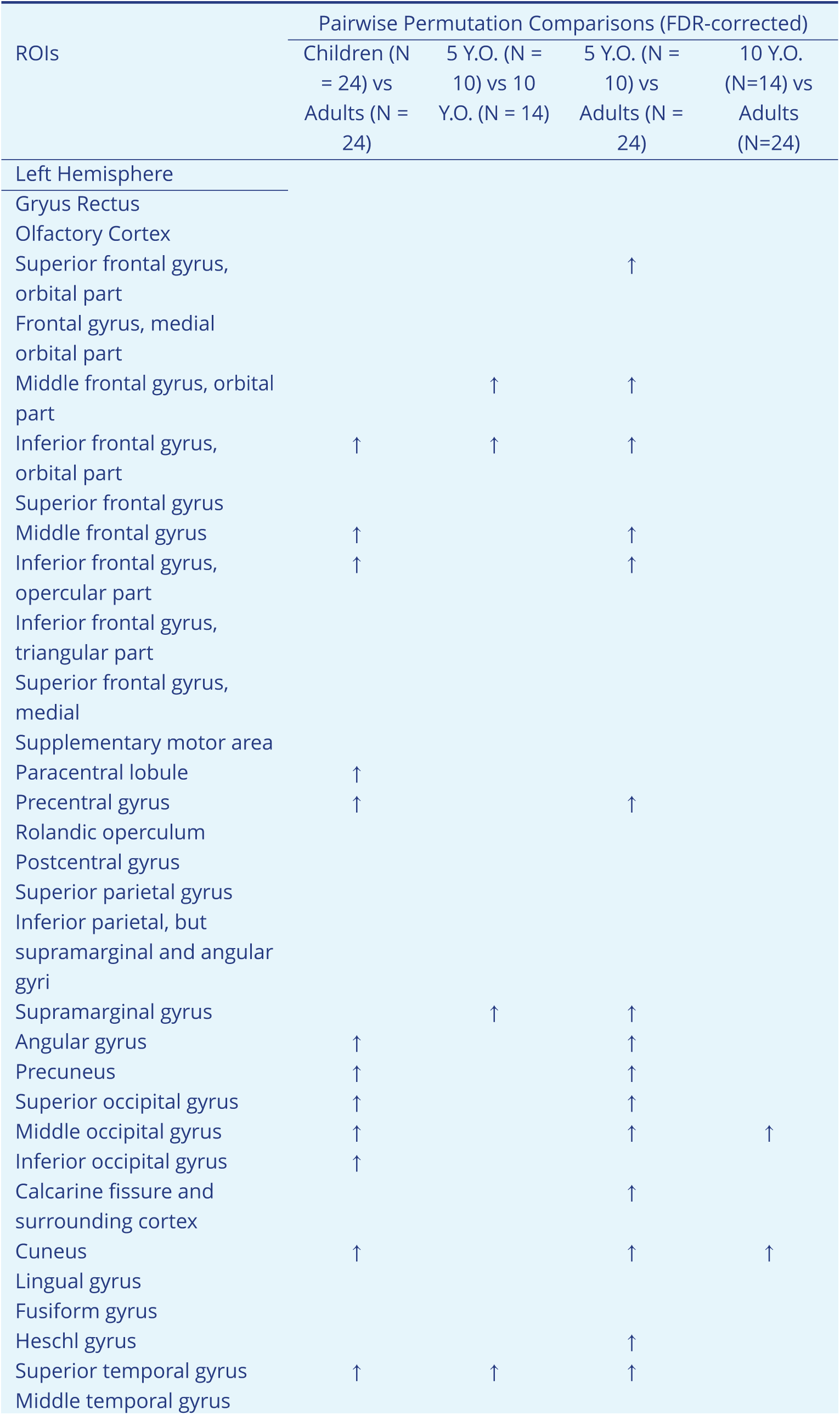
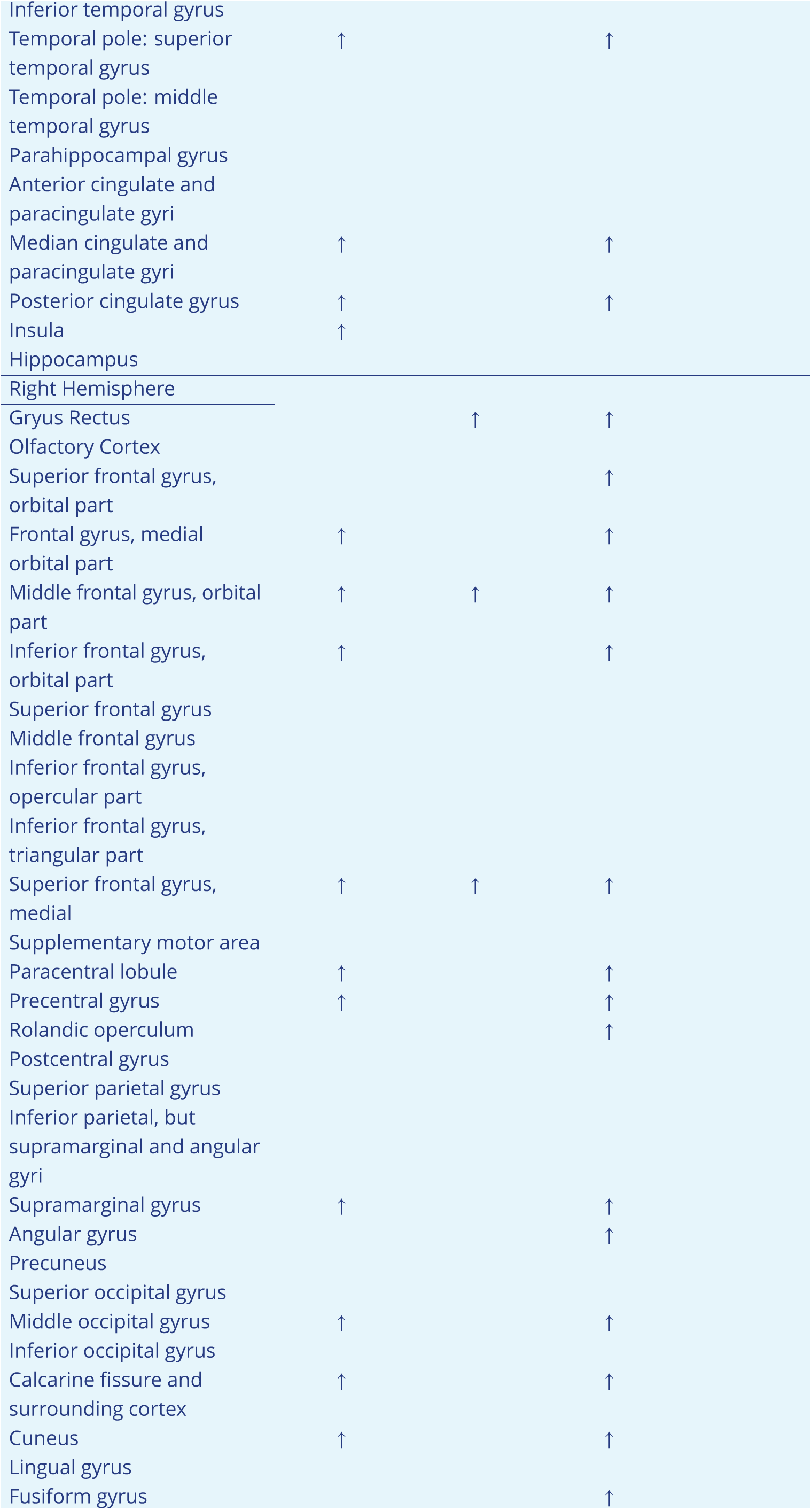
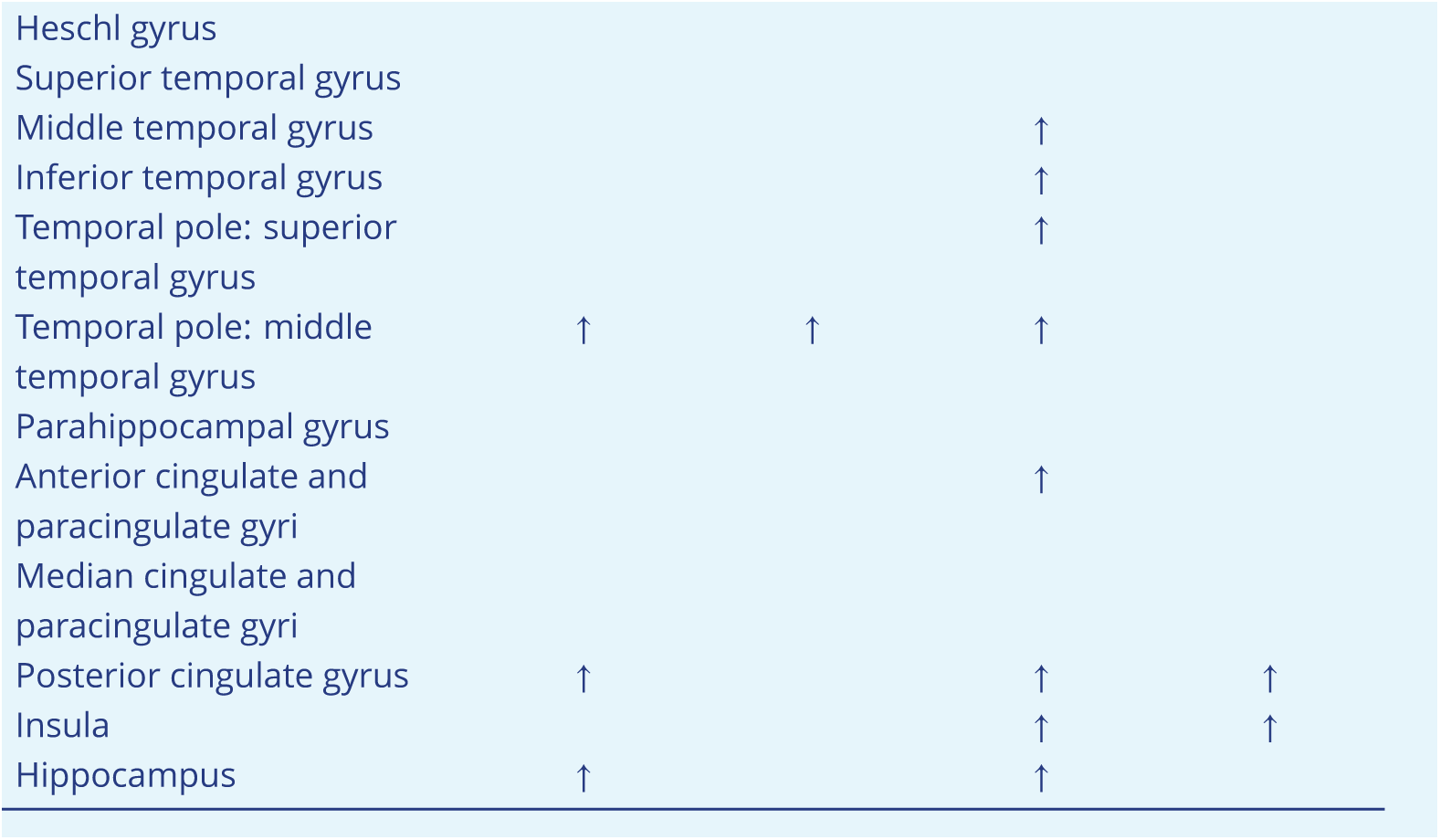
Regions of interest (ROIs) that manifest significant Eccentricity differences between groups in the low gamma band.

| ROIs | Pairwise Permutation Comparisons (FDR-corrected) |  |  |  |
| --- | --- | --- | --- | --- |
|  | Children (N = 24) vs Adults (N = 24) | 5 Y.O. (N = 10) vs 10 Y.O. (N = 14) | 5 Y.O. (N = 10) vs Adults (N = 24) | 10 Y.O. (N=14) vs Adults (N=24) |
| Left Hemisphere |  |  |  |  |
| Gryus Rectus |  |  |  |  |
| Olfactory Cortex |  |  |  |  |
| Superior frontal gyrus, orbital part |  |  | ↑ |  |
| Frontal gyrus, medial orbital part |  |  |  |  |
| Middle frontal gyrus, orbital part |  | ↑ | ↑ |  |
| Inferior frontal gyrus, orbital part | ↑ | ↑ | ↑ |  |
| Superior frontal gyrus |  |  |  |  |
| Middle frontal gyrus | ↑ |  | ↑ |  |
| Inferior frontal gyrus, opercular part | ↑ |  | ↑ |  |
| Inferior frontal gyrus, triangular part |  |  |  |  |
| Superior frontal gyrus, medial |  |  |  |  |
| Supplementary motor area |  |  |  |  |
| Paracentral lobule | ↑ |  |  |  |
| Precentral gyrus | ↑ |  | ↑ |  |
| Rolandic operculum |  |  |  |  |
| Postcentral gyrus |  |  |  |  |
| Superior parietal gyrus |  |  |  |  |
| Inferior parietal, but supramarginal and angular gyri |  |  |  |  |
| Supramarginal gyrus |  | ↑ | ↑ |  |
| Angular gyrus | ↑ |  | ↑ |  |
| Precuneus | ↑ |  | ↑ |  |
| Superior occipital gyrus | ↑ |  | ↑ |  |
| Middle occipital gyrus | ↑ |  | ↑ | ↑ |
| Inferior occipital gyrus | ↑ |  |  |  |
| Calcarine fissure and surrounding cortex |  |  | ↑ |  |
| Cuneus | ↑ |  | ↑ | ↑ |
| Lingual gyrus |  |  |  |  |
| Fusiform gyrus |  |  |  |  |
| Heschl gyrus |  |  | ↑ |  |
| Superior temporal gyrus | ↑ | ↑ | ↑ |  |
| Middle temporal gyrus |  |  |  |  |

|  |  |  |  |
| --- | --- | --- | --- |
| Inferior temporal gyrus |  |  |  |
| Temporal pole: superior temporal gyrus | ↑ |  | ↑ |
| Temporal pole: middle temporal gyrus |  |  |  |
| Parahippocampal gyrus |  |  |  |
| Anterior cingulate and paracingulate gyri |  |  |  |
| Median cingulate and paracingulate gyri | ↑ |  | ↑ |
| Posterior cingulate gyrus | ↑ |  | ↑ |
| Insula | ↑ |  |  |
| Hippocampus |  |  |  |
| Right Hemisphere |  |  |  |
| Gyrus Rectus |  | ↑ | ↑ |
| Olfactory Cortex |  |  |  |
| Superior frontal gyrus, orbital part |  |  | ↑ |
| Frontal gyrus, medial orbital part | ↑ |  | ↑ |
| Middle frontal gyrus, orbital part | ↑ | ↑ | ↑ |
| Inferior frontal gyrus, orbital part | ↑ |  | ↑ |
| Superior frontal gyrus |  |  |  |
| Middle frontal gyrus |  |  |  |
| Inferior frontal gyrus, opercular part |  |  |  |
| Inferior frontal gyrus, triangular part |  |  |  |
| Superior frontal gyrus, medial | ↑ | ↑ | ↑ |
| Supplementary motor area |  |  |  |
| Paracentral lobule | ↑ |  | ↑ |
| Precentral gyrus | ↑ |  | ↑ |
| Rolandic operculum |  |  | ↑ |
| Postcentral gyrus |  |  |  |
| Superior parietal gyrus |  |  |  |
| Inferior parietal, but supramarginal and angular gyri |  |  |  |
| Supramarginal gyrus | ↑ |  | ↑ |
| Angular gyrus |  |  | ↑ |
| Precuneus |  |  |  |
| Superior occipital gyrus |  |  |  |
| Middle occipital gyrus | ↑ |  | ↑ |
| Inferior occipital gyrus |  |  |  |
| Calcarine fissure and surrounding cortex | ↑ |  | ↑ |
| Cuneus | ↑ |  | ↑ |
| Lingual gyrus |  |  |  |
| Fusiform gyrus |  |  | ↑ |
| Heschl gyrus |  |  |  |  |
| Superior temporal gyrus |  |  |  |  |
| Middle temporal gyrus |  |  | ↑ |  |
| Inferior temporal gyrus |  |  | ↑ |  |
| Temporal pole: superior temporal gyrus |  |  | ↑ |  |
| Temporal pole: middle temporal gyrus | ↑ | ↑ | ↑ |  |
| Parahippocampal gyrus |  |  |  |  |
| Anterior cingulate and paracingulate gyri |  |  | ↑ |  |
| Median cingulate and paracingulate gyri |  |  |  |  |
| Posterior cingulate gyrus | ↑ |  | ↑ | ↑ |
| Insula |  |  | ↑ | ↑ |
| Hippocampus | ↑ |  | ↑ |  |

